# DEPP: Deep Learning Enables Extending Species Trees using Single Genes

**DOI:** 10.1101/2021.01.22.427808

**Authors:** Yueyu Jiang, Metin Balaban, Qiyun Zhu, Siavash Mirarab

## Abstract

Placing new sequences onto reference phylogenies is increasingly used for analyzing environmental samples, especially microbiomes. However, existing placement methods have a fundamental limitation: they assume that query sequences have evolved using specific models directly on the reference phylogeny. Thus, they can place single-gene data (e.g., 16S rRNA amplicons) onto their own gene tree. This practice is a proxy for a more ambitious goal: extending a (genome-wide) species tree given data from individual genes. No algorithm currently addresses this challenging problem. Here, we introduce Deep-learning Enabled Phylogenetic Placement (DEPP), an algorithm that learns to extend species trees using single genes without pre-specified models. We show that DEPP updates the multi-locus microbial tree-of-life with single genes with high accuracy. We further demonstrate that DEPP can achieve the long-standing goal of combining 16S and metagenomic data onto a single tree, enabling community structure analyses that were previously impossible and producing robust patterns.

## Introduction

In recent years, phylogenetic inference has found widespread application in identifying organisms that make up a biological sample (Hebert et al., 2003; Seifert et al., 2007; Munch et al., 2008). Microbiome analyses often rely on phylogenetic analyses to identify species present in an environment sampled using the 16S rRNA gene amplicon sequencing or whole metagenome shotgun sequencing data (Handelsman, 2004; Langille et al., 2013; Sunagawa et al., 2013; Matsen, 2015; N.-p. Nguyen et al., 2014; Truong et al., 2015; Asnicar et al., 2020). Using the phylogenetic context, we can identify species even when exact matches to the reference datasets are not present. Similarly, outside microbiome analyses, identifying new and known species using (meta)barcoding and genome skimming data rely on phylogenetic analyses (Kress et al., 2009; QUICKE et al., 2012; Ballesteros and Hormiga, 2018; Bohmann et al., 2020).

In these high-throughput applications, phylogenetic placement – adding a new sequence onto an existing reference tree – is sufficient, and the more challenging *de novo* reconstruction is neither necessary nor always more accurate (Janssen et al., 2018). Phylogenetic placement has a long history of method development (Felsenstein, 1981; Desper and Gascuel, 2002; Mirarab, N. Nguyen, et al., 2012; Matsen et al., 2010; Berger et al., 2011; Barbera et al., 2019; Balaban, Sarmashghi, et al., 2020; Rabiee and Mirarab, 2020). However, these algorithms add sequences from a single gene family onto a tree showing its evolutionary history (i.e., the gene tree). Phylogenetic relationships change across the genome (Maddison, 1997; Degnan and Salter, 2005) due to processes such as horizontal gene transfer (HGT) (Ochman et al., 2000), and accounting for such discordance is a subject of much recent method development (Warnow, 2017). Given data from individual genes, it must be assumed that they have evolved on a gene tree, not the species tree. Thus, existing methods place on gene trees but use the gene tree as a proxy for the species tree, hoping that their differences are not consequential. For example, species identification using marker genes such as 16S or COI (e.g., Konstantinidis and Tiedje, 2005; Zaneveld et al., 2010) implicitly assumes that the gene tree is close enough to the species tree to allow accurate identification of the *species* by placement on the gene tree. More recently, Rabiee and Mirarab, 2020 enabled inserting a genome onto a species tree by minimizing quartet distance, but this approach requires genome-wide data.

Users of phylogenetic placement often face a question: given query sequence data from a single gene (or a handful of genes), is it possible to place the query onto the species tree (a goal we name *discordant placement*). The correct position of a query on the species tree is not always determinable from a single gene. Nevertheless, the gene tree is related to the species tree, and gene data have *some* information about the correct placement on the species tree, giving us hope that discordant placement can be achieved with sufficient if imperfect accuracy.

The ability to extend a species tree using single-gene data would be tremendously useful in studying ecology. Microbiome analyses are split between metagenomic and 16S data, which remain mostly disconnected. Accurate discordant placement methods would enable researchers to add 16S samples onto species trees and treat them as if they were metagenomic samples, albeit with less signal and more uncertainty. Thus, it would become possible to combine 16S and metagenomic data (Fig. 1) in downstream analyses such as sample differentiation (Matsen IV et al., 2013) using methods such as UniFrac (Lozupone and Knight, 2005) and taxonomic profiling. In particular, the ability to add *all* types of data,including 16S, metagenome-assembled genomes (MAGs), and marker genes on the same backbone tree using the same algorithm will be helpful in ensuring consistency for downstream applications. For example, we often have a large number of 16S samples and a smaller number of metagenomic samples (presumably with better differentiation ability) available to study the impact of microbiome on a phenotype. Inserting all samples onto the same tree will enable probing the associations between composition of samples and phenotypes using a unified analysis. Similarly, the ability to place eukaryotic species sampled only through their barcode genes (e.g., COI) on species trees will greatly benefit ecology research.

**Figure 1:**
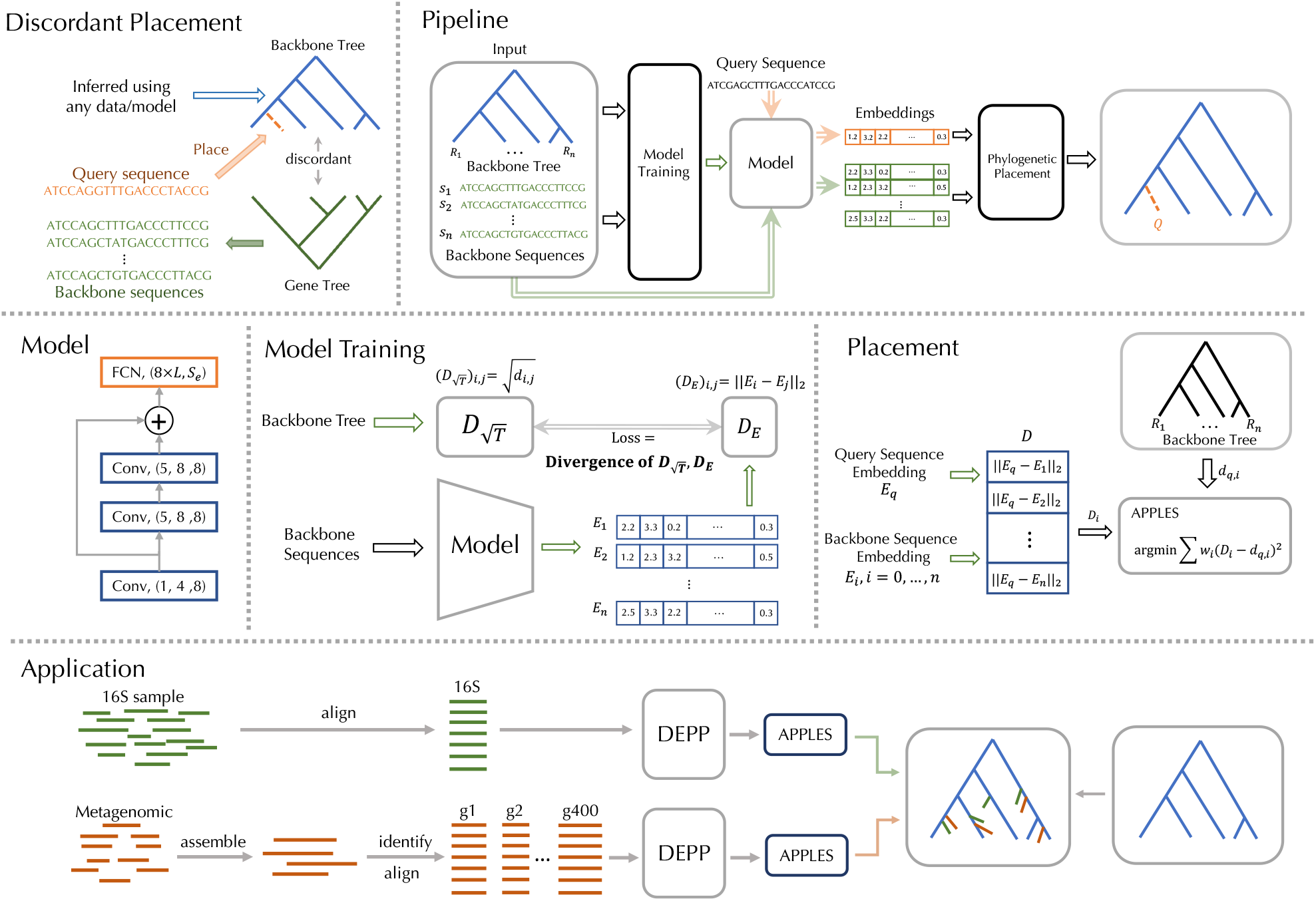
Overview of the method. Top: an illustration of discordant placement and the over-all process. A neural network model is trained from a given reference tree, however inferred, and reference sequences labeled by the leaves of the tree. The model maps sequences to embeddings in *k* dimensions. Once sequences are embedded, phylogenetic placement is performed using distances between embeddings of the query and reference species. Middle: the structure of the CNN model and both training and placement processes. The training process minimizes the difference between Euclidean distances in the ℝ_*k*_ space and the square root of tree distances across all pairs of species. The placement process computes Euclidean distances between embeddings of the query and references. The resulting distance vector is placed using weighted least square error minimization (APPLES). Bottom: The process for combining 16S and metagenomic samples using DEPP.

How can we approach the discordant placement problem? One solution is to misuse the existing methods and simply assume the species tree and the gene tree are the same. However, it is unclear how these methods are expected to behave when the sequence data has not directly evolved on the tree. The standard phylogenetics methods are model-based: they assume data has evolved on a tree according to a model of sequence evolution (e.g., GTR Tavaré, 1986) and infer the tree using maximum likelihood, Bayesian sampling, or model-corrected distances (Warnow, 2017). While this paradigm has enjoyed much success, its reliance on explicit models can make it less accurate in the face of deviations from assumptions of the model (Jermiin et al., 2020; Naser-Khdour et al., 2019; Lartillot et al., 2007; Sullivan and Swofford, 2001; Halpern and Bruno, 1998). Using existing methods for the discordant placement problem would lead to a misspecified model, which can result in low accuracy, as we show. We suggest that instead of relying on the dominant paradigm, discordant placement can be solved using machine learning.

As early as the 1990s, researchers attempted phylogenetic inference using general purpose machine learning models, such as neural networks. Dopazo and Carazo, 1997 formulated the problem as unsupervised learning and designed a neural network that reflected the tree shape. More recently, the success of deep neural networks (DNNs) in solving other challenging problems has motivated efforts to adopt DNNs in phylogenetics. Zou et al., 2020a and Suvorov et al., 2020 have used a semi-supervised approach with two steps: for every four species (a quartet), classify the input data to one of the three possible quartet topologies, then combine these quartet trees. This formulation raises a question: where are we to find labeled training data in the high volume needed by DNNs? Both papers turn to simulations for an answer: use complex models to simulate data on known trees, from which we then train the model. The point of these methods is to use complex generative models that can be sampled but do not avail themselves to scalable inference. However, learning from simulated data runs the risk of missing relevant parts of the huge parameter space and model misspecification. As Zaharias et al., 2021 recently showed, these methods can have lower accuracy than standard methods in careful benchmarking.

Phylogenetic placement offers a way to use general purpose models without simulations. Given a reference tree, however computed, and sequence data which are a function of the tree, we can use the reference data to train a machine learning model (such as a DNN) that can place a query sequence onto the reference tree. The reference tree may be a species tree inferred using large numbers of genes, using complex models, and perhaps after spending much computational resources. Such reference data are increasingly available. For example, several comprehensive trees were published in the past year on tens of thousands of microbial species (Zhu, Mai, et al., 2019; Parks et al., 2020; Asnicar et al., 2020) using 120 to 400 genes, with analyses that took *>* 200, 000 hours of CPU and GPU time in one case. These available trees are excellent candidates for providing the training data.

Why would we need to turn to black-box machine learning models for discordant placement? Note that discordant placement only poses a weak connection between sequence data and the tree: the sequence data should be a function of the tree. In contrast, traditional models require that sequence data have evolved *directly* on the tree according to a specific Markov model, which is a much stronger assumption. A compelling reason to use machine learning is that it provides ways to learn general functions (here from sequences to positions on trees). By avoiding explicit mechanistic models, machine learning has the potential to build general models that map sequences onto trees even when the tree and the sequences are not fully compatible, enabling placement onto species trees using single genes. Such a model would, in effect, simultaneously place sequences onto a gene tree and *reconcile* (Doyon et al., 2011) the differences with the species tree.

In this paper, we introduce the Deep-learning Enabled Phylogenetic Placement (DEPP) framework (Fig. 1). Given a reference tree, inferred in any way, and some sequence data labeled by leaves of the tree but potentially evolved on a tree incongruent with the reference tree, DEPP learns a neural network to embed sequences in a high dimensional Euclidean space, such that pairwise distances in the new space correspond to the square root of tree distances. Given such a model, the placement of new sequences can proceed by computing the embedding, computing distances, and using distance-based phylogenetic placement.

## Materials & Methods

### Discordant phylogenetic placement

The standard phylogenetic placement takes as input a *reference* tree *T*, its associated sequences *S*, and a query sequence *q*. The output is the best placement of *q* on *T*, which consists of a specific position of a particular edge of *T* and the length of a new terminal branch. In contrast to the standard placement, we assume the relationship between the reference tree and sequences is indirect. Thus, *S* (typically sequences from a single gene) does not directly evolve on *T* (typically a species tree) but is influenced by *T*. Also, *T* can be inferred from any source of data (not just *S*). We define the discordant phylogenetic placement as the problem of finding the best placement of the query on *T* using *S despite* disagreements between *S* and *T*. What “best placement” depends on the context. When the discordance between data and the reference tree is because *T* is a species tree but the sequence data come from a single gene, we define the ideal placement as the true placement of the species on the species tree. This definition is meant to enable the application laid out before; namely, updating the species tree and identifying samples using a single gene. If this problem could be solved completely accurately, we could build species trees using single genes, and we could fully identify samples using their marker genes. Alternative definition of “best” could be imagined. For example, we could seek the place of queries that would minimize the distance between the updated tree and the true gene tree; such definitions may be appropriate for other applications.

### Background on Distance-based Estimation

Let *T* denote a rooted phylogenetic tree on a set of *n* taxa 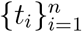 represented as leaves and each branch labeled with its length. The tree *T* defines a distance matrix where each entry *d*_*ij*_ ∈ ℝ^+^ is the path length between leaves *i* and *j*. A distance matrix may or may not be equal to that of some tree, but when it does match a tree, it matches a *unique* tree and is called additive (Buneman, 1974). Assume we are given a set of sequences *S*, and each *s*_*i*_ ∈ *S* corresponds to a leaf *t*_*i*_. Computing distance between sequences produces a sequence distance matrix. These distances can converge to additivity if computed under the correct statistical model. For example, under the Jukes and Cantor, 1969 (JC) model, 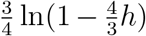 asymptotically converges to additivity where *h* is the hamming distance between sequences. Note that in discordant phylogenetic placement, traditional methods for obtaining distances from *S* would not match *d*_*ij*_ as the sequences have not evolved on the reference tree directly.

### Placement as supervised learning

We approach discordant placement using supervised learning (Fig. 1). The training data are *T* and *S* and our model is a convolutional neural network (CNN). We use a distance-based approach and express the training data as {((*s*_*i*_, *s*_*j*_), *d*_*ij*_)} where *s*_*i*_, *s*_*j*_ ∈ *S* are for pairs of reference sequences and *d*_*ij*_ is the distance between taxa *i* and *j* on the tree *T*. Due to the discordance, which leads to model misspecification, such using JC or similar distances may not be accurate. Instead, we use machine learning to compute distances. In addition, since real datasets almost always include missing data, we use machine learning to also reconstruct the missing parts of a sequence to obtain more accurate distances.

### Learning Objective

To compute distances, we build a CNN that embeds sequences in the ℝ^*k*^ space; we then use distances between embeddings as estimates of phylogenetic distances. The use of embeddings enjoys a theoretical justification. As Vienne et al., 2012 showed and Layer and Rhodes, 2017 elaborated, for any tree *T*, there *exists* a collection of points 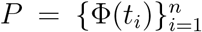 in the ℝ^*n*−1^ Euclidean space such that the distance between the points Φ(*t*_*i*_) and Φ(*t*_*j*_) equals to 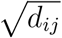. Thus, if sequences are a function of the tree, there must exist an embedding that corresponds to the tree, and we use machine learning to find an embedding that minimizes the divergence between embedding distances and (the square root of) distances on the given reference tree.

More precisely, we treat the reference tree *T* and sequences *S* as training data and seek a model that maximizes the match of Euclidean distances between embeddings and the square root of phylogenetic distances in the reference tree (i.e.,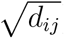). The square root is to match the theory by Vienne et al., 2012 and Layer and Rhodes, 2017. To make our goal precise, we need to define a measure of matrix similarity. While any metric can be used (and measures such as log-determinant divergence have shown promise (e.g., Xie et al., 2018)), here, we simply use mean squared error, seeking:

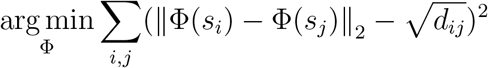

where Φ : {*A, C, G, T*}^*L*^ → ℝ^*k*^ is an embedding of sequences in the Euclidean space, and *d*_*ij*_ give pairwise path distances in the reference tree *T*. While we focus on nucleotides here, a similar formulation can be used for amino acid sequences or any type of character data. Note that the estimated distance of *i* and *j* is (‖Φ(*s*_*i*_) − Φ(*s*_*j*_)‖_2_)^2^.

Motivated by strong evidence in distance-based phylogenetics that weighting down long distances improves accuracy (Gascuel, 2000; Desper and Gascuel, 2002; Balaban, Sarmashghi, et al., 2020), we define a weighted version of the objective function:

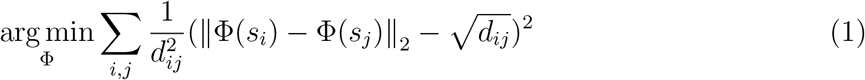

which follows the Fitch and Margoliash, 1967 (FM) principle. Note that other weighting schemes (e.g., Beyer et al., 1974; Desper and Gascuel, 2002) can be used and will be tested.

### Embedding size

The Layer and Rhodes, 2017 (LR) formulation requires *n* − 1 dimensions, which introduces some challenges. According to the theory, the number of dimensions needs to increase by one after inserting the query. Our supervised learning formulation does not allow that (the embedding size is fixed after training). Thus, there is no guarantee that the embeddings remain correct after addition, even if they are before addition. However, we note that, in LR embeddings, adding a leaf would require simply dividing one of the *n*−1 dimensions into two dimensions, leaving the rest of the embeddings intact. Thus, one can hope that having one less dimension has a minimal practical impact. More broadly, for large *n*, training models with *n*-dimensional embedding is impractical. Thus, we often set *k < n* − 1, and the gap can be more than an order of magnitude for some of our tests described below. In practice, we use a rule of thumb to select the default *k* (which the user can change), setting 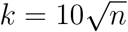, rounded to the nearest power of 2 from below (i.e., 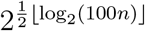).

### Model structure and training

#### CNN

We use a convolutional neural network (Fig. 1). Nucleotide sequences are encoded using 4-bit one-hot binary vectors; we refer to each bit as a channel. Gaps can be encoded as all zeros (DEPP version ≤ 0.1.13) or by setting all four channels to 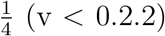. Moreover, as detailed below, we can use a separate model to guess the best values to represent a gap (v ≥0.2.0). To accommodate reference species that have multiple copies of a gene, we change the encoding so that instead of a binary vector, it includes the frequencies of the nucleotide characters among the gene copies. These encodings provide the input “features” that are processed through three linear convolutional layers, each followed by a nonlinear layer and a fully-connected linear layer.

As detailed in Appendix C, a convolutional layer applies a set of parameterized kernels by convolving them across its input (i.e., using the dot product of the kernel entries and the input). Convolutional layers are usually used as feature extractors, and multiple layers are used to detect high-level abstraction from the input. Here, we use them to enable the model to go beyond the traditional i.i.d models of sequence evolution and capture *k*-mer signatures. The first convolutional layer takes as input *L* features, each encoded as four channels, and outputs an 8 × *L* matrix by applying a kernel size of 1 (but applied to all four input channels). The next two convolutional layers each have a kernel size of 5 (i.e., operating on 5-mers). The input of each layer is padded with zeros on both sides so that the output has the same length as the input. The input to the second convolutional layer is added to the output of the third convolutional layer, forming a residual block, which is a cornerstone of deep learning (He et al., 2016). Using residual blocks can help solve the vanishing gradient problem, which is why they are commonly used, including for sequence data analyses (Killoran et al., 2017; Zou et al., 2020b). To enable the model to capture nonlinear relations, after each convolutional layer, we use a nonlinear layer built using the continuously differentiable exponential linear unit, which has performed well in other contexts (Barron, 2017). The last layers are fully connected, taking features from outputs of convolutional layers and producing the final embeddings or the probability vectors. This layer aggregates the signal from convolutional layers. Each activation in the output is connected to all the inputs, and the output is a weighted sum of all the inputs.

#### Handling missing data

Multiple sequence alignments almost always include gaps, which may represent missing data or indels. While indels may represent real signal, here, just like traditional maximum likelihood phylogenetic models, we treat all gaps as missing data. As mentioned earlier, we can deal with missing data by leaving the one-hot encoding ambiguous (e.g., setting all four channels to 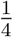). An alternative is to amputate the missing data using. Motivated by the BERT (Devlin et al., 2018) model used extensively in NLP, DEPP includes a reconstruction neural network to guess the best encoding for missing data (implemented since v0.2.0?). The input of the model is a sequence with missing data (gaps) encoded as a one-hot vector, and the output is the reconstruction of the sequence where the sites with gaps are probability vectors inferred by the model. During training, we randomly select sites and label them as gaps to provide supervised signal for learning the reconstruction objective. This learning objective is the Kullback-Leibler divergence between the one-hot encoding of the letter and the output probability vector:

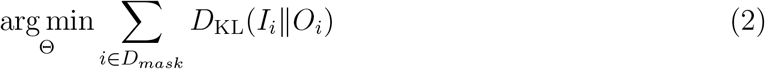

where *D*_*mask*_ are randomly chosen masked sites, and *I*_*i*_ and *O*_*i*_ are the one-hot encoding of the letter and the output probability vector for the site *i*. The reconstruction model, which consists of three convolutional layers and one fully-connected layer, is trained separately from the DEPP encoder. At the time of testing/placement, a query sequence is first run through the reconstruction model to fill in the gaps, and the output of the reconstruction network is fed into the DEPP encoder to generate embeddings.

#### Training

We trained the model with Eq. 1 as the loss function for DEPP encoder and Eq. 2 as the loss function for reconstruction network using the stochastic gradient descent algorithm RMSProp, which divides the gradient by a running average of its recent magnitude to speed up training (Tieleman and Hinton, 2012). The batch size is fixed to 32. We check the training loss every 50 epochs and stop the training when the value of the loss function fails to decrease in two consecutive checks. The model with the optimal objective function value is chosen.

### Placement

Once the CNN model is trained, we use it to map a given query sequence *q* to a vector of distances *D*_1_ … *D*_*n*_. For datasets with missing data (gaps), we compute two sets of distances,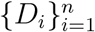 and 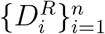, using the models with and without gap reconstruction, respectively. The final distances is set to the weighted sum of the distances, i.e. 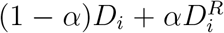 where *α* is the proportion of the sites with gaps in the query sequences. The weighted sum is used to reduce the impact of reconstructed bases (which are guessed, as opposed to being observed) on the final distance and will be empirically tested. Given these distances, we then place *q* onto *T* using distance-based placement (Balaban, Sarmashghi, et al., 2020), which uses dynamic programming to find the placement with the minimum 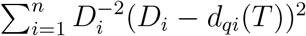 where *d*_*qi*_(*T*) represents the tree-based distance between the query and each taxon *i* (Fig. 1).

### Uncertainty calculation

In principle, bootstrapping, the dominant method used in phylogenetics, can be used to estimate uncertainty around distances and thus placements. However, bootstrapping assumes i.i.d models, and convolutional networks like DEPP do not treat input as i.i.d. Moreover, bootstrapping would require retraining our model on each replicate bootstrap, which we do not afford. Instead, we measure the uncertainty using a subsampling procedure based on solid grounds from the non parametric support estimation literature. For each query, we randomly select *m* sites and mask them as gaps. The masked sequences are input to the pretrained DEPP model, which produces a distance vector corresponding to the distances from the query to backbone species. We repeat this step *r* times and get *r* distance vectors 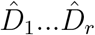. A correction is then applied to the *r* distance vectors as

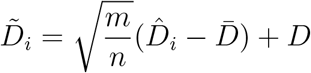

where 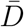 is the average over 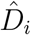, *D* is the distance vector corresponds to the sequence with no site masked, and *n* is the length of the sequence. We use 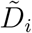 for placement and get *r* placements for each query, which are then used to calculate support of the placements by counting the number of times each edge is chosen. In our experiment, we choose *r* = 200 and set *m* = *n*/log^0.1^(*n*).

The 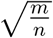 term adjusts for the increased variance of estimates obtained from fewer data points, and is justified as follows. For a statistically consistent estimator 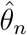 of a parameter *θ* based on *n* data points, if there is some rate of convergence *τ*_*n*_ such that 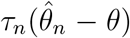 weakly converges to *some* distribution as *n* → ∞ (assumption 2.2.1 of Politis et al., 1999), then, under forgiving conditions, the distribution of 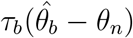 converges to that of 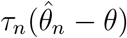 as *n* → ∞ as long as *b* → ∞ and *b/n* → 0 (see Theorem 2.2.1 in Politis et al., 1999). While the choice of *τ*_*n*_ is not obvious in general, by central theorem limit, for any estimator that can be described as the sum of independent random variables, 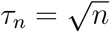 is the correct choice. Motivated by this observation, we set 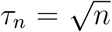 which gives us the 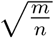 correction term, noting that we have no proof that the rate of convergence of our estimator is proportional to 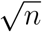 (or for that matter, that our estimator is consistent). We will evaluate the support values empirically.

### DEPP implementation Details

We implemented DEPP using PyTorch and treeswift python packages (Moshiri, 2020) and trained the models on 2080Ti NVIDIA GPUs. The embedding size *k* is set to 128 for datasets with 200 taxon and 512 for larger datasets (including the real WoL dataset). Other hyperparameters are fixed to their defaults (Table S2) unless otherwise specified. DEPP is trained on the reference tree and is used to compute distances that are then fed to APPLES-II (Balaban, Jiang, et al., 2021). APPLES-II is used identically to APPLES-II+JC (see below). Branch lengths of the backbone tree provided to DEPP are reestimated using RAxML-8 (Stamatakis, 2014) under the GTR+CAT model. Given more than one gene, DEPP has two options: concatenating genes or computing a summary of distances. For each query, we can compute the distance between a query and backbone species *j* according to each of the *N* genes, obtaining 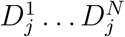 (ignoring missing genes). We summarize all 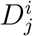 values by setting *D*_*j*_ to the average of all 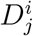 values that fall between 25 and 75 percentiles of all 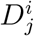 values (to remove the impact of outlier genes).

### Methods compared

We compared DEPP v.0.2.2 (unless otherwise specified) to three methods.

#### EPA-ng

(Barbera et al., 2019) This maximum likelihood method is widely used but is *not* designed for discordant placement. Nevertheless, we test it for placing gene data on the species tree, with branch lengths re-estimated using RAxML-ng, under the GTR + G model.

#### INSTRAL

(Rabiee and Mirarab, 2020) This method updates a species tree given input gene trees (already updated to include the query) and accounts for discordance by maximizing the quartet score. Here, input gene trees are inferred using FastTree-II (Price et al., 2010) or RAxML. Results with FastTree or RAxML give similar results (Fig. S16), so in the main paper we use the results with input tree built by FastTree-II. While INSTRAL does account for discordance, it requires *at least two* genes and is designed for cases with many genes.

#### APPLES+JC

We use APPLES in its default settings where it computes distances using Jukes Cantor (JC) model (Jukes and Cantor, 1969), chosen because Balaban, Sarmashghi, et al., 2020 found no evidence that more complex models improve accuracy. The branch lengths of reference trees are re-estimated based on the JC model using RAxML-ng (Kozlov et al., 2019). APPLES also includes two options *d*_*m*_ and *b*_*m*_ to set *w*_*qi*_ = 0 (i.e., ignore distances) for at most *n* − *b*_*m*_ reference taxa per query when *δ*_*qi*_ *> d*_*m*_. We use *d*_*m*_ = 0, *b*_*m*_ = 5 for all datasets.

### Datasets

#### Simulated dataset

For studying the gene tree and species tree discordance due to incomplete lineage sorting (ILS), we use a published simulated dataset (Mirarab and Warnow, 2015), with 200 ingroup species and gene trees that are discordant with the species tree due to ILS. The dataset contains model conditions corresponding to high, medium, and low ILS, each with 50 replicates. We arbitrarily selected the first 2^0^, … 2^5^ genes for each replicate.

We also simulated a second dataset that consists of gene trees and species trees discordant due to horizontal gene transfer (HGT) in addition to low levels of ILS. We used Simphy 1.0 (Mallo et al., 2016) to simulate 10 replicates, each with 10,000 ingroup species and 500 genes. Species trees are simulated using 10^8^-generation birth-death process (Kendall, 1948) with birth and death rates set to 5 × 10^−7^ and 4.167 × 10^−7^, respectively. This would lead to low levels of ILS —average normalized RF distance between species and gene trees due to ILS is 0.03. In addition to ILS, each gene goes through HGT at a rate drawn from lognormal distribution with *µ* ∼ 𝒩 (−18, 0.4), *s*^2^ = 0.75. Simphy uses a HGT model that reduces the probability of transfer proportionally to the distance between source and recipient. The combined effect of ILS and HGT leads to 0.44 (normalized RF) median gene tree discordance. Given true gene trees simulated using Simphy, we simulate alignments with length ranging between 231 and 2054 using Indelible (Fletcher and Yang, 2009). Since training a model on 10,000 species takes considerable computational resources, we then pick five genes per each replicate. To do so, we ordered the genes by the number of species that are horizontally transferred and pick 50th, 150th, 250th, 350th, and 450th genes in that the ordered gene list to ensure our test cases include genes with different level of HGT.

#### Web-of-life (WoL) marker genes

Zhu, Mai, et al., 2019 built a species tree of 10,575 prokaryotic genomes using ASTRAL from 381 marker genes and computed mutation unit branch lengths using 100 sites randomly selected from each of the 381 marker genes. We categorized the marker genes into three equal-sized groups based on the rank of quartet-distance (Sand et al., 2013) between their gene tree and the species tree. For each discordance category, we selected the 10 most commonly present genes among all species (Table S3). The mean gene tree discordance with the species tree, measured using the quartet distance, is 0.18, 0.34, 0.50 in low, medium, and high discordant groups. Gene alignments are available from Zhu, Mai, et al., 2019.

#### WoL rRNA genes

16S and 5S rRNA genes were predicted using RNAmmer (Lagesen et al., 2007) and aligned using UPP (N.-p. D. Nguyen et al., 2015). In genomes with multiple copies of 16/5S, we train and test DEPP on all copies and re-estimate backbone tree branch lengths for APPLES+JC using an arbitrary copy. Due to their wide usage in microbiome analyses, we also perform analyses with three regions of 16S commonly used in amplicon sequencing: V3+V4 (≈ 400bp), V4 (100bp), and V4 (150bp). We removed from 16S datasets any predicted sequence output by RNAmmer that was shorter than half of the average sequence length (removing less than 1% of species; see Table S4).

#### Traveler’s Diarrhea microbiomes

Quality-controlled 16S rRNA gene amplicons and manually curated MAGs were derived from a study by Zhu, Dupont, et al., 2018, which identified novel pathogenic profiles from the fecal samples of 22 Traveler’s Diarrhea (TD) patients as compared with seven healthy traveler (HT) controls. The 16S rRNA amplicon sequence variants (ASVs) were generated using Deblur from QIIME 2 and are 250bp long. The 381 marker genes were identified using PhyloPhlAn on the translated protein sequences inferred by Prodigal from the contigs included in each MAG. This protocol is identical to that used in the WoL study.

### Evaluation procedure and leave-out experiments

To ensure query sets (i.e., testing data) are separate from the training data, we removed 5% of species (10 and 500 for simulations with 200 and 10,000 backbone taxon and ≈ 445 for real data) from the species tree to obtain reference trees. We did not re-estimate species trees after removing queries. These left-out species are used as the query. The reference tree is the true species tree for the simulated dataset and the ASTRAL tree from the WoL dataset (Zhu, Mai, et al., 2019). For the simulated dataset, branch lengths of the backbone species tree are estimated using sites randomly selected from the genes we used in the experiments (32 genes for ILS data and 5 genes for HGT data) with each gene providing 500 sites. For WoL dataset, branch lengths of the species tree are available from the original study (estimated under GTR+Γ from 100 randomly selected sites from 381 marker genes). Training is done using DEPP v0.2.2. For testing, each query taxon is placed independently, and the result is compared against the full reference tree before pruning the query (i.e., the true tree for simulations and the ASTRAL tree for WoL). The error metric we report is the number of edges between the position on the reference tree and the inferred placement. In total, we have 8,934 and 25,000 test cases for the ILS simulated and HGT simulated data respectively and 14,616 test cases for the WoL dataset. INSTRAL fails in 66/9000 tests; we exclude these cases for all the methods.

In addition, we categorize test cases by their level of ILS, level of HGT, and phylogenetic signal. We compute the level of ILS by measuring the Robinson and Foulds, 1981 (RF) distance between true gene trees and the species tree. The phylogenetic signal is a function of many factors, including sequence length, tree height, and the rate of evolution. Here, to quantify the lack of signal, we use the RF distance between true gene trees and those estimated using FastTree-II (Price et al., 2010). These two measurements are per backbone tree. In contrast, we measure HGT levels on a per query basis by inspecting species close to the query species in the true gene tree and their placement in the species tree. Specifically, for the five nearest species in the gene tree to the query *q* (denote them by *N*_5_), we compute the sum of their path length (number of branches) to *q* in the species tree. Note that this sum can never be less than 17, which is the value obtained if *N*_5_ are the five closest leaves to *q* in the species tree and the topology is identical and balanced. We measure HGT as the average path length of *N*_5_ above 17; i.e., 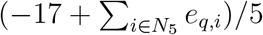 where *e*_*q,i*_ is the number of branches from the query *q* to species *i* in the species tree. Thus, 0 means the query is placed in a similar context in the gene tree and thus there is no HGT *close* to that leaf, while a high value indicates that the species close to the query in the gene tree are far away from it in the true species tree, indicating HGT.

### Case study on Traveler’s Diarrhea

For ASV placement, a single model is trained using DEPP v.0.2.2 with the reference tree set to the WoL species tree (backbone tree) and backbone sequences coming from V4 region of 16S (≈250bp). The trained model is applied on the ASV in the Traveler’s Diarrhea dataset to calculate the distance matrix between the sequences in the studied dataset and the backbone sequences. We removed one gene (p0150), where all backbone sequences were gapped for at least half of the sites. Finally, we train a model using DEPP v.0.2.2 on the remaining sequences, giving us 380 models in total. We release and maintain these reference DEPP models for public use (see Data and Code Availability); v1.0.0 of the database is used in these analyses.

For placing MAGs, first, we used UPP to extend the alignments of all 380 marker genes to include the markers identified in the TD dataset. Given these alignments, we used DEPP and the 380 trained models to compute distances between query marker genes and backbone WoL species, resulting in 380 distance matrices. We then use then distance-summary option of DEPP (i.e. mean distance in the interquartile range) to summarize all distances and use APPLES-II for placement.

To compare pairs of samples, we use weighted UniFrac (Lozupone and Knight, 2005; McDonald et al., 2018). A feature table containing the frequency of ASVs or the number of reads matching a MAG in each sample is available. We use the feature table and the placement tree to calculate weighted UniFrac between each pair of the samples using QIIME 2 (Bolyen et al., 2019). We then use the PERMANOVA (Anderson, 2001), as implemented in QIIME 2, to compare the HT group and the TD group (the number of permutation is set to be 10^6^ − 1). To visualize the correlation between samples, we apply Principal Coordinates Analysis (PCoA) on the weighted UniFrac distance matrix using QIIME-2 (P. Legendre and L. Legendre, 2012; Halko et al., 2010), picking the top 3 coordinates for visualization.

We calculated a *MAG coverage* metric for each sample to represent the proportion of sequencing data covered by a MAG. It equals (∑_*i*∈*M*_ *L*_*i*_*C*_*i*_)*/*(∑ _*i*∈*N*_ *L*_*i*_*C*_*i*_), where *M* is the set of contigs constituting MAGs, *N* is the set of all contigs in a metagenome assembly, *L* is the length (bp) of a contig, *C* is the coverage of a contig as determined by the average number of times each nucleotide of the contig is included in any sequencing read recruited to the contig.

## Results

### Evaluation on simulated datasets

#### DEPP training and parameter sensitivity

We start by evaluating DEPP on simulated datasets, testing the ability to train the CNN model in reasonable times. As the training epochs advance, the loss function (1) drops rapidly and stabilizes after around 500 epochs in a typical case (Fig. S1). Here, training, which is a one-time process for each reference tree, finished in around 20 minutes for the 200-taxon dataset and 260 minutes for 10000-taxon dataset, on a machine with one 2080Ti NVIDIA GPU and 8 CPU cores; placement of 1000 queries took 4 seconds for the 200-taxon and 30 seconds for the 10000-taxon datasets using a single CPU core.

DEPP is mostly robust to weighing schema, with all four schemes tested resulting in statistically indistinguishable performance (Table S2). Models with more parameters, i.e., deeper network or larger embeddding size, tend to have better performance but also longer training time. For example, for the 200-taxon tree, the time for training a model with one residual block is around 15 minutes while this number goes up to 20 minutes when the model has 5 residual blocks. Reducing the number of residual blocks from five to one or reducing the embedding size to 32 reduce the accuracy significantly (Table S2). In our final model, we use five residual blocks for backbone tree with 200 taxon and one residual block for the rest of dataset to trade-off performance and training time. Note that while the preliminary results motivated the choice of default settings (Table S2) used in the rest of analyses, we did not select the optimal settings for this dataset and have not tested various settings on other datasets (thus, hyperparameters are not overfit to the data). The use of the gap reconstruction model dramatically improves accuracy when the query has 40% or more gaps, and the use of the weighted approach results in further reductions in error (Fig. S17).

#### Comparison to other methods

We now compare accuracy of DEPP to distance-based APPLES-II (Balaban, Jiang, et al., 2021) used with the standard Jukes Cantor (JC) model, maximum likelihood method EPA-ng (Barbera et al., 2019), and the quartet-based discordant-aware method INSTRAL (Rabiee and Mirarab, 2020). Note that APPLES-JC and EPA-ng are not designed for discordant placement using a single gene, and INSTRAL is designed only for datasets with many genes (at least two but ideally many more). However, since no existing method is designed for discordant placement, we had to compare it to these existing methods.

#### ILS discordance

On the 200-taxon dataset, DEPP is comparable to EPA-ng and outperforms APPLES+JC and INSTRAL when given a single gene (Fig. 2ab). While DEPP and EPA-ng have similar average error rates overall, DEPP has fewer cases with very high errors (Fig. 2a). When the gene tree discordance level is medium (or low), DEPP, EPA-ng and APPLES+JC have similar average error, which is as low as 1.5 edges (or 0.9 edge) on average but DEPP has a shorter error tail (Fig 2a). DEPP has the lowest mean error among the methods when discordance is high (3.38 edges for DEPP versus 3.50 for EPA-ng and 4.31 for APPLES+JC). For context, random placements on these species trees would give a placement error of 14 edges on average (empirically computed). Furthermore, DEPP outperforms other methods in difficult cases when the phylogenetic signal is weak (Fig. 2b).

**Figure 2:**
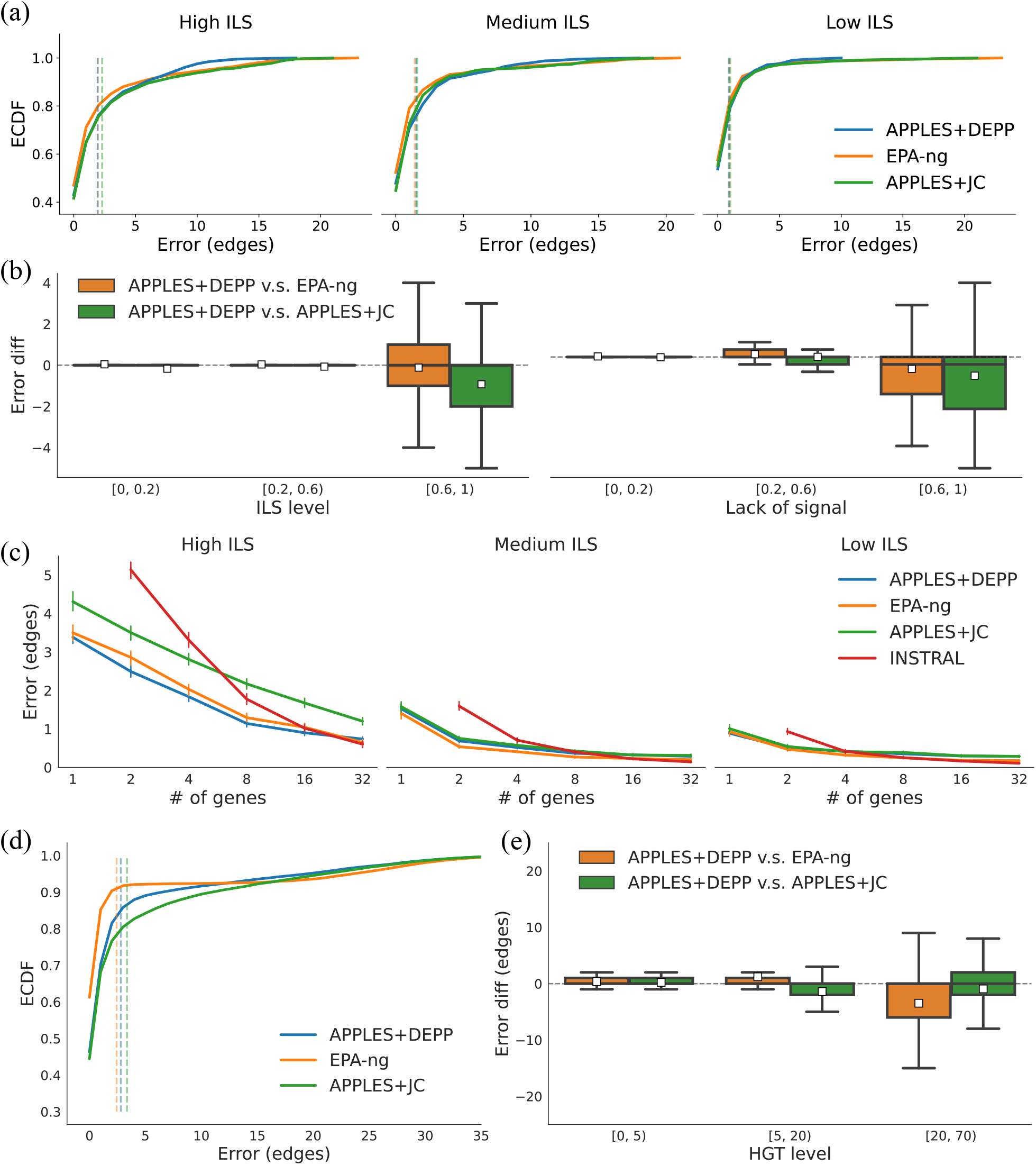
Results on simulated datasets. a) Empirical cumulative distribution function (ECDF) of the placement error on a single gene for high, medium, and low discordance (69% 34%, and 21% mean RF). INSTRAL needs at least 2 genes. b) Mean and standard error of placement error versus the number of genes on ILS data. c) Sensitivity to gene properties on ILS data. Error comparison between DEPP and other method using a single gene on (left) different level of true gene tree discordance (RF distance between true gene trees and the species tree) and (right) different level of gene signal missing (RF distance between true gene trees and estimated gene trees) combining all discordance levels. y-axis: the error of DEPP minus error an alternative method. White squares: mean error difference. d) Empirical cumulative distribution function (ECDF) of the placement error on HGT data. e) Error comparison between DEPP and other method on different level of HGT. HGT is measured by 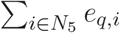, where *N*_5_ is the 5 closest species on the gene tree and *e*_*q,n*_ is the number of branches between queries and species *i* on species tree.

All methods experience a sharp rise in the error when the phylogenetic signal weakens (Fig. S5a) or discordance increases (Fig. S5bc). According to an ANOVA test (Table S1), both gene tree discordance and signal have a significant impact on the placement error (*p*−value*<* 10^−20^). However, these two factors combined explain only around 15% of the variance in error for DEPP and EPA-ng.

As the number of concatenated genes increases, unsurprisingly, the mean errors of all methods reduce (Fig. 2c). Computing per-gene distances and summarizing them instead of concatenating them increases accuracy under some conditions but reduces accuracy under others (Fig. S2). DEPP, EPA-ng as well as APPLES+JC are more accurate than INSTRAL for low numbers of genes (≤ 4) but not for more genes. In fact, as the number of genes increases to 32, INSTRAL starts to have the best accuracy, a result consistent with the theory as INSTRAL is statistically consistent under ILS.

We further examine the example cases where DEPP performs well or poorly. We compare the ML phylogenetic distances computed using RAxML on the true species tree versus distances computed by DEPP for low and high error cases (Fig. 3). Both high and low error cases seem to result in unbiased distances computed in the training set; however, high error examples have higher variance. The high variance can be attributed to a lack of signal: two identical sequences in the gene may belong to different parts of the species tree, a problem that the model cannot overcome. At the time of testing (e.g., for a query), distances are systematically overestimated in cases with high error, and a large range of values are estimated for pairs with equal ML distances. In some cases, DEPP assigns small distances to some reference species that have high distances to the query (Fig. 3; bottom left).

**Figure 3:**
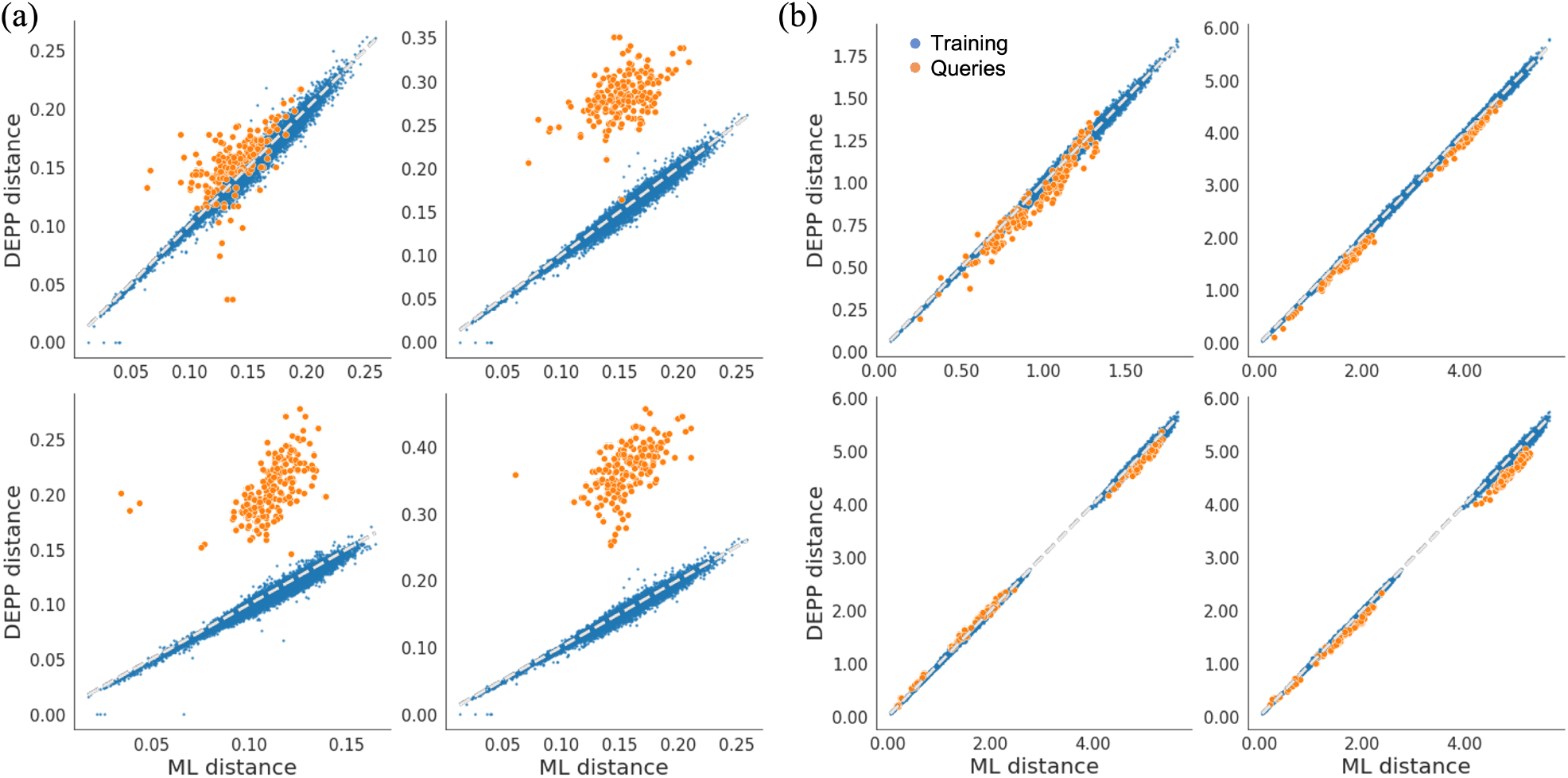
Examples of phylogenetic distance calculation using DEPP. Phylogenetic distance on the true species tree (in units of substitution per site) versus the distance calculated by DEPP for (a) examples with high error (11 edges of error) and (b) examples with zero error. Dashed gray line: identity line. High errors correspond to long terminal and short internal branches; see Fig. S3 for trees.

Examining example trees shows that these cases of high error tend to correspond to novel query taxa; i.e., those on long branches on sparsely sampled clades (Fig. S3). Queries with higher terminal branch lengths lead to higher error (Fig. S4); however, it appears that the shortest of branches also have a higher error, perhaps because distinguishing very similar taxa requires strong signal. Similarly, queries with a large clade as the sister tend to be more difficult for DEPP (Fig. S4). Trees that lead to high error tend to have long terminal branches and short branches close to the root, a condition that corresponds to rapid radiations; in contrast, easy cases are those with shorter terminal branches and long branches closer to the root (Fig. S3).

#### HGT discordance

On the 10000-taxon dataset, which includes HGT, EPA-ng has the best overall accuracy, with DEPP coming as a close second (mean error: 2.43 and 2.80, respectively). However, these average performances mask larger differences as HGT levels change. Breaking down the dataset by the HGT level per query taxon, we observe that DEPP has slightly worse accuracy than EPA-ng for the queries with low or medium HGT level but better accuracy with high levels of HGT. DEPP has the largest advantage with the most challenging cases when gene sequences moved far away; e.g., error for DEPP is 3.5 edges better than EPA-ng and 0.9 edges better than APPLES+JC on average with the highest level of HGT.

#### The real WoL dataset

We then tested DEPP on the real WoL dataset using 30 marker genes, pre-selected to represent of the range of discordance among all 381 genes, in addition to 16S and 5S. We tested DEPP, EPA-ng and APPLES+JC using both novel queries (left-out) and observed queries (training data). Despite the size of the datasets, neither DEPP training nor placement was prohibitively slow. For example, on the 16S gene with 7407 species, training takes 240 minutes and uses 5GB of memory using a 2080Ti NVIDIA GPU. Placement of 800 queries on the backbone takes less than 40 seconds using a single core.

Given large backbone trees such as WoL, a query sequence will have a considerable chance of matching a species present in the reference tree. Under such conditions, DEPP has better performance over all the genes compared with EPA-ng and APPLES+JC (Fig. S6). DEPP has close to perfect accuracy for all genes except the multi-copy genes 16S and 5S (Figs. 4a and S6). For these genes, the error slightly increases; DEPP finds the optimal placement in 91% (16S) and 73% (5S) of cases but, on average, has an error of 0.32 (16S) and 1.13 (5S) edges. Patterns of error do not change whether we train DEPP using the weighting scheme described before or by simply selecting an arbitrary copy (Fig. S7). While APPLES+JC and EPA-ng also had low error levels for many genes, they had considerable error levels on the training datasets for eight and three genes, respectively (Fig. S6).

**Figure 4:**
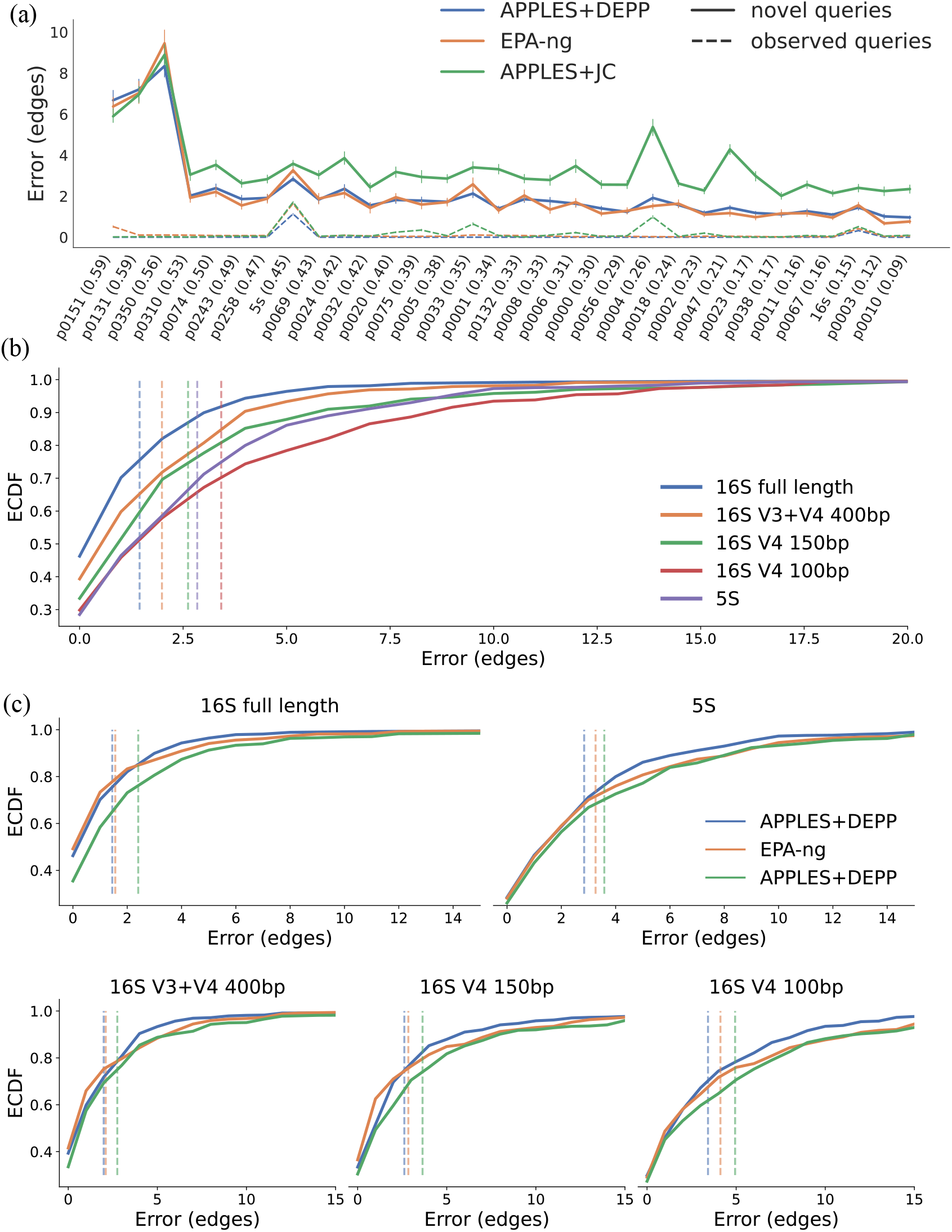
Results on the WoL dataset. a) Mean and standard error of placement error for APPLES+JC, EPA-ng and DEPP, applied to novel queries and known queries, on the species tree from WoL dataset, which is treated as the ground truth. The x-axis shows genes, ordered by their quartet distance to the reference species tree (shown parenthetically). For novel queries, the placement is performed in a leave-out fashion. For Empirical Cumulative Distribution (ECDF) of the placement error, see Fig. S8. b) ECDF of placement error on the 16S gene with full length or individual regions of 16S used in amplicon sequencing. V3+V4 region is ≈ 400bp. Dashed lines show the mean error. C) Methods comparison on rRNA genes. We cut the x-axis at 15 for better visualization; see Fig. S11 for the full range.

In the most interesting case, when the query sequences are novel (i.e., are not in the training set), both DEPP and EPA-ng greatly outperform APPLES+JC (Figs. 4a). On average, the placement error of DEPP (2.17 edges) and EPA-ng (2.15 edges) is much lower than APPLES+JC (3.34 edges). Moreover, EPA-ng and DEPP have low error about the same number of times (respectively, 89% and 88% of DEPP and EPA-ng placements have four edges or less error), DEPP is less often far away from the optimal placement. For example, on average, the maximum error of DEPP for each gene is 7 edges lower than EPA-ng; or, the placement error of EPA-ng is larger than 15 edges in 3.1% of cases compared to 2.4% for DEPP. Thus, just like the simulated dataset, DEPP has fewer cases of complete failure.

Across all 32 genes, in 85% of tests, DEPP and EPA-ng finds a placement within 3 edges of the optimal placement; for the full-length 16S gene, this value is 91% for DEPP and 87% for EPA-ng. A random placement on a tree with 10,575 leaves is, on average, 26 edges away from the optimal placement. Our results are also consistent with using 16S as a marker gene, which among the 32 genes had one of the lowest mean error rates (1.43 edges). Finally, note that similar to simulated datasets, accuracy of DEPP is a function of the accuracy of its distances. Calculated distances have very little bias and high variance where DEPP works well (Fig. S9) but high variance and bias when it works poorly (Fig. S10).

Going beyond a single gene, given 50 randomly selected genes, DEPP used with distance summary strategy (see Materials & Methods) was able to place within three branches of the optimal placement in 94% of cases, with a mean error of only 0.98 edges (Fig. S12). Accuracy slightly degrades if we concatenate genes instead of summarizing distances among them (e.g., mean error increases to 1.6 edges). Thus, DEPP can not only place using single genes, but it can also place with high accuracy given data from multiple genes.

Testing performance on rRNA genes 16S and 5S, average error for DEPP is lower than EPAng or APPLES+JC for both genes (Fig 4c). Mean error for DEPP over all rRNA data is 2.48 edges compared to 2.81 for EPA-ng and 3.47 for APPLES+JC. For full-length 16S, DEPP and EPA-ng find the optimal placements about the same number of times; however DEPP is more often *close* to the optimal placements (e.g., 91% of queries are within three edges for DEPP compared to 87% for EPA-ng and 80% for APPLES+JC). When given short amplicon-length sequences, the advantage of DEPP over alternatives becomes more substantial. For example, given 100bp amplicons from the V4 region, DEPP has an average error of 3.4 edges while the average error of EPA-ng and APPLES+JC are 4.12 and 4.94 edges, respectively. While 3.4 edges of error may sound high, note that insertion of a 100bp read into a *species tree* is clearly a difficult task. Comparing performance on different rRNA genes, for 16S data, longer sequences give better performance for all the methods (Fig. 4). Interestingly, 5S sequences (which are ≈100bp) have better accuracy than 16S V4 region sequences with the similar length, possibly indicating that 5S carries more phylogenetic signal or less discordance with the species tree.

#### Support accuracy

We test our method of measuring support on 30 marker genes of WoL data versus the support values generated by EPA-ng. Support values are vastly different between EPA-ng and DEPP. While EPA-ng tends to produce 100% support for a single placement for most queries, DEPP generate far more placements, most with low support (Fig 5a). Across all 12,727 queries, EPA-ng generates only 14,355 placements versus 163,110 produced by DEPP. DEPP estimates full support for a single placement for only 9.8% of queries whereas EPA-NG produces full support for 85% of queries. Because of its high confidence in its unique placement, no threshold of EPA-NG support can produce low levels of false positive detection (Fig 5b), in contrast to DEPP, which can produce FPRs close to zero. Comparing FPR and recall, DEPP can achieve the same recall level as EPA-NG with far lower levels of FPR (Fig 5b).

**Figure 5:**
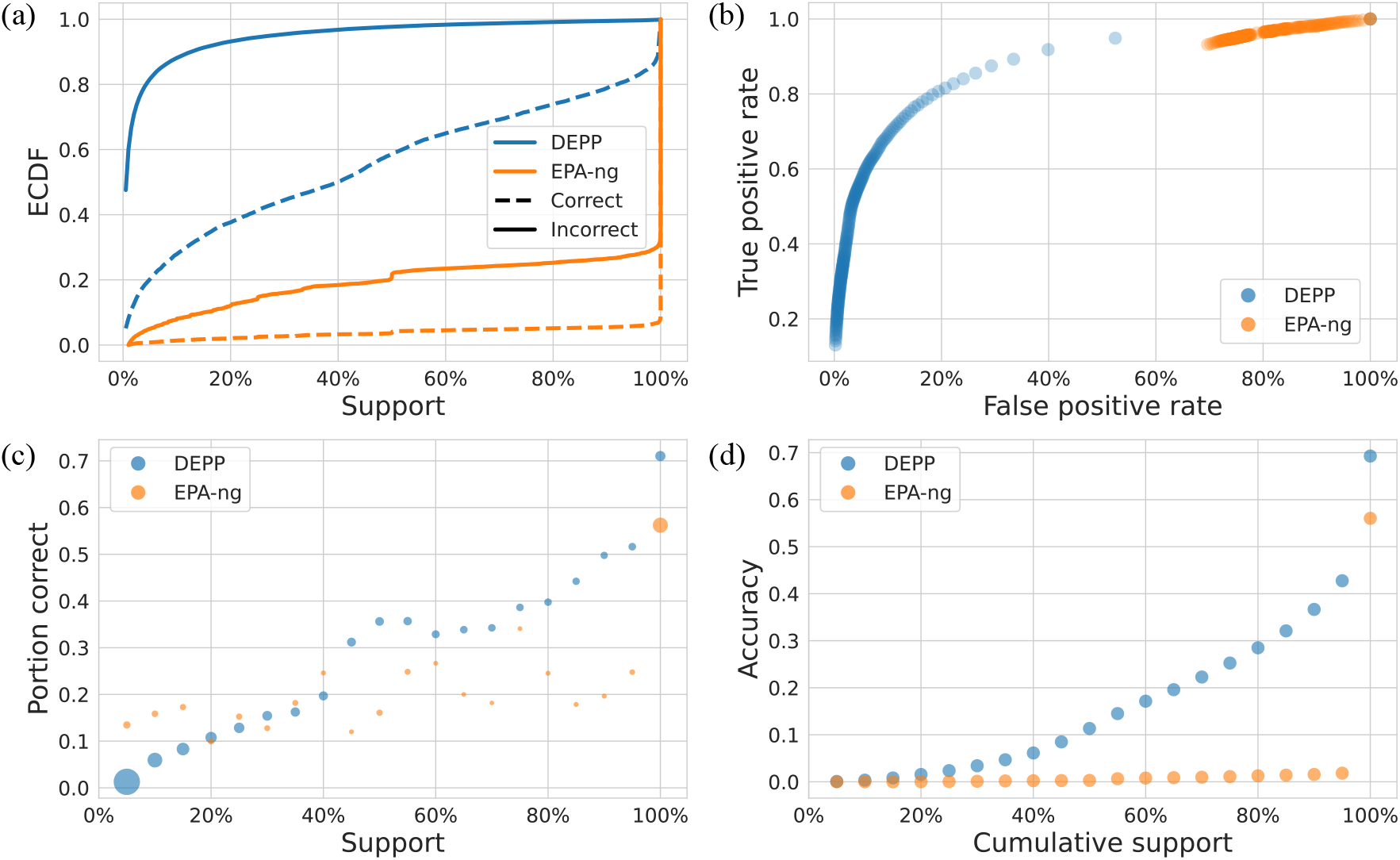
Support of the placements. a) Empirical cumulative distribution function (ECDF) of the supports. b) Receiver operating characteristic (ROC) curve. TP, TN, FP, FN is defined by correct placement with support above the threshold, incorrect placement with support below the threshold, incorrect placement with support above the threshold, correct placement with support below the threshold respectively (support threshold step size: 0.5%). c) Correlation between support and the correctness of the placements (area of each dot is proportional to the number of placements it contains). d) Correlation between support and placement accuracy. x-axis: cumulative support of the top placements (placements with highest supports); y-axis: proportion of the cases when the correct placements are among the top placements. These figures are using 163110 placements for DEPP and 14355 placements for EPA-ng in total across 12727 queries on 30 marker genes of the WoL dataset.

Both EPA-NG and DEPP support values tend to be higher in distribution for correct placements than incorrect placements (Fig 5ac). However, DEPP support values show a much larger gap between correct and incorrect branches and are more predictive of accuracy compared to EPA-ng (Fig 5ac). Both methods clearly over-estimate support so that even branches with 100% support are often not incorrect; for example, only 66%, 71% and 76% of the DEPP placements with support ≥ 0.9, ≥ 0.95 and ≥ 0.99 are correct. However, the over-estimation problem is worse for EPA-ng, which gives high support in the vast majority of cases (Fig 5cd). Overall, 62% of wrong placements with EPA-ng have 100% support, compared with only 0.2% for DEPP.

### Case study on combined 16S rRNA and shotgun metagenomic data

We next studied how adding the the metagenome-assembled genomes (MAGs) and 16S rRNA amplicon sequence variants (ASVs) onto the same tree enables new analyses. We used the dataset by Zhu, Dupont, et al., 2018 with gut microbiomes from seven healthy controls (HT) and 22 patients with Traveler’s Diarrhea (TD), all of whom were sampled using both 16S amplicon sequencing and metagenomics with available MAGs and ASVs. We added the ASVs and MAGs onto the same WoL backbone tree using DEPP (see Materials & Methods) obtaining two placement profiles for each subject. We compare pairs of profiles using the weighted UniFrac (Lozupone and Knight, 2005) distance.

The UniFrac distances of the MAG profile of a sample to the ASV profiles of other samples were higher on average than its distance to the ASV profile of the same sample in all except one case (Fig. 6a). In nine samples, the MAG profile had a lower distance to its own ASV than *any* other sample. The intra-sample distances between the ASV and MAG profiles substantially reduced as the MAGs represented a larger proportion of the sequencing data of each sample (Fig. 6a; *p*-value = 0.001), whereas distances across samples slightly increase (*p*-value = 0.02). As a result, the gap between inter- and intra-sample distances grew substantially with higher MAG coverage (Fig. 6a).

**Figure 6:**
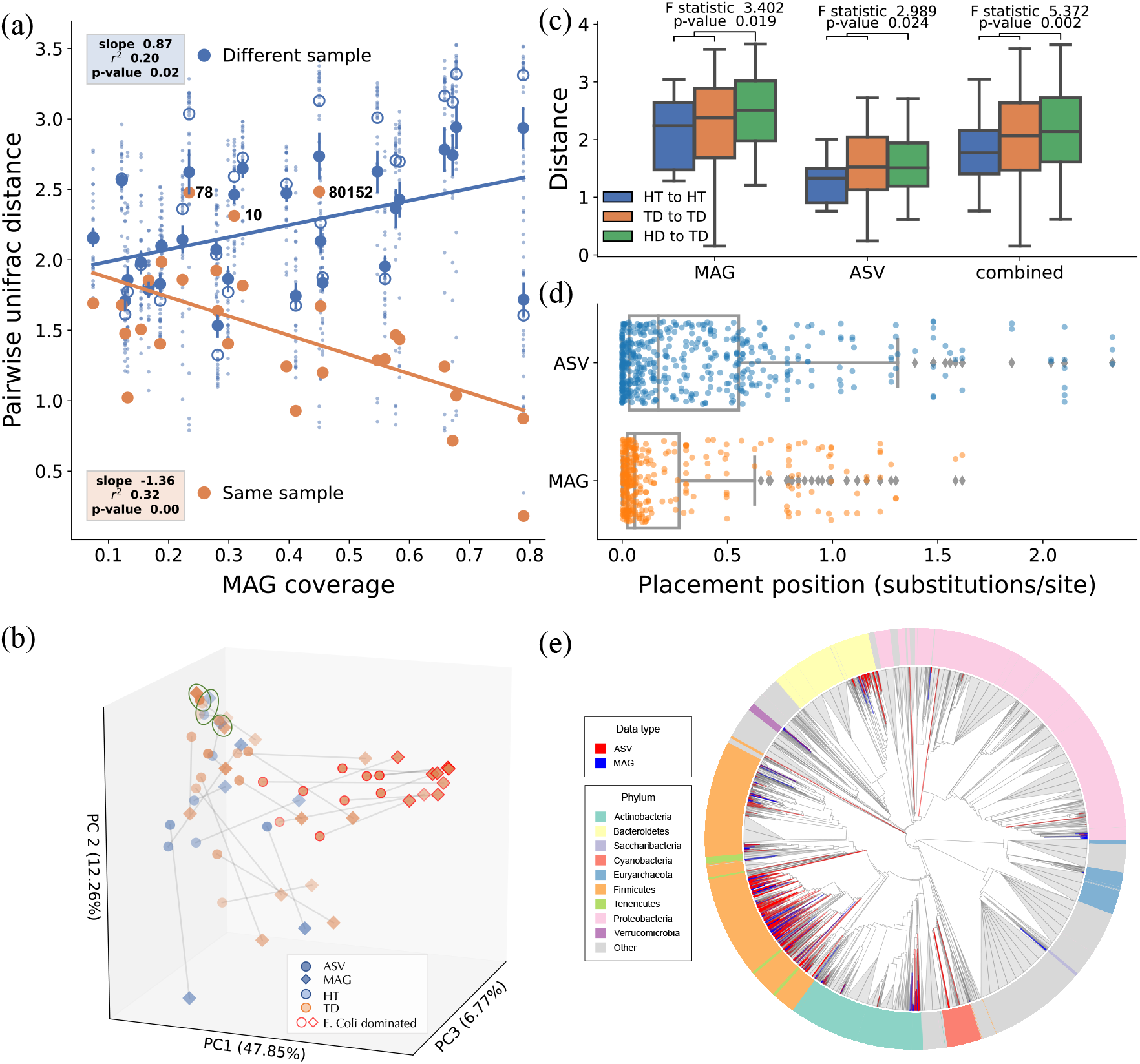
Combined MAG and ASV results on the TD dataset. a) For every sample, the UniFrac distance (*y*-axis) is shown between its MAG placements and ASV placements of the same sample (orange) and ASV placements of other samples (blue) versus the MAG coverage (proportion of sequencing data covered by the MAGs). For inter-sample comparisons, we show all pairs of comparisons (small dots), the mean and standard error (large sold dots and bars), and the median (empty circles). b) PCoA analyses of combined ASV and MAG placements based on weighted UniFrac Distances. Each sample is represented with two connected dots, one for MAG and one for ASV. Examples of samples where ASV and MAG have very low distances are highlighted in green. Nine samples dominated by *E. coli* are highlighted with a red border. c) Distribution of UniFrac distances among pairs of samples within HT or TD group and across the groups, using MAGs, ASVs or the two data combined. F-statistic and p-values are calculated using the PERMANOVA test. d) Distance of the placement position (geometric mean of tMRCA of the species in the closest sibling clade to the query) using ASV and MAG data; each dot represents one bin or ASV. e) The ASV and MAG placements on the WoL phylogenetic tree (grey: backbone taxa).

Placements on a single tree enabled us to visualize all ASV- and MAG-informed community structures using a unified Principal Coordinates Analysis (PCoA) (Fig. 6b). This results show that for some samples, ASV and MAG placements indicate extremely similar community structures while for others, there is a substantial disagreement between the two. The intra-sample distances in the PCoA plots tend to be shorter for high coverage MAGs (Fig. S13a).

MAGs and ASVs can both distinguish healthy and diseased samples (*p*-values: 0.019 and 0.024 using the standard PERMANOVA method), but MAGs provide a higher statistical power that implies a larger effect size (Fig. 6c). While the median ASV distances between pairs of TD samples are similar to distances between TD and HD samples, both intra-group median MAG distances are lower than inter-group MAG median. Furthermore, combining two type of data, 16S and MAG, which is enabled by DEPP, provides a large separation with much larger F statistics and increased statistical significant (*p*-values: 0.002) (Fig. 6c).

Despite the substantial agreement between MAG and ASV placements, there are also differences. For three samples (78, 10, 80152), the ASV/MAG agreement is low compared to the background distance levels. Zhu, Dupont, et al., 2018 characterized these samples as suffering from the co-infection of multiple Enterobacteriaceae organisms (*Escherichia, Enterobacter, Klebsiella*, and *Citrobacter*), which increased the challenge of accurately binning contigs from these closely related microbes – a possible explanation for the relatively low congruence. In addition, there are several groups that are found by MAGs or ASVs but not the other (Figs. 6e, S15). The MAG placements include a clade representing Saccharibacteria which is not represented in the ASV placements. This group of bacteria is classified under the candidate phyla radiation (CPR), which has distinct physiological and genetic characteristics from all remaining bacteria. Commonly used 16S rRNA primers have reduced sensitivity in capturing the Saccharibacteria group (Castelle and Banfield, 2018) that, according to Zhu, Dupont, et al., 2018, may be responsible for the disease status. On the other hand, the ASV placements include several clades under Cyanobacteria not found by MAGs. These may represent the commonly seen contamination from chloroplasts of dietary plants in 16S analyses (Di Rienzi et al., 2013), a problem that does not afflict metagenomic assembly.

Beyond the detected groups, branch lengths also reveal interesting patterns. While MAGs tend to be placed close to the tips, a majority of ASV placements are deep in the tree (Fig. 6d). These more basal placements of ASVs are consistent with the lower phylogenetic signal included in short sequences, which can lead to less specific characterizations. Shorter terminal branches highlight the advantage of MAGs versus 16S rRNA amplicons in understanding microbiome compositions.

We observed substantial correlation (*r*^2^: 0.57; *p*-value *<* 10^−5^) between the Faith’s phylogenetic (alpha) diversity computed using ASV and MAG placements (Fig. S14). These strong but imperfect correlations once again show the two sources of data capture similar but subtly different patterns. Overall, due to the more basal placements discussed earlier, alpha diversity measured using ASV tends to be higher. Several low alpha diversity cases are related to nine TD samples reported by Zhu, Dupont, et al., 2018 to be dominated by *E. coli*. The first axis in the combined PCoA analysis clearly separates these samples from others using MAGs, but the separation is less strong using ASVs, a pattern observed if PCoA analyses are performed separately for ASV and MAGs (Fig. S13bc).

## Discussion

We introduced a deep learning approach for extending an existing phylogenetic tree without a need for pre-specified models of sequence evolution or gene tree discordance. Our approach learns how to add new taxa by capturing patterns in an existing reference set. Thus, it uses the backbone alignment and tree to learn a model that maps sequences onto the tree. This automatic learning of the model eliminates the need for assuming rigid models. Given a correct sequence evolution model and no discordance, we see no reason machine learning should be more accurate than traditional phylogenetics. The power of DEPP is in its ability to learn from data without knowing the model. For example, in our simulations where the GTR+Gamma model of sequence evolution and MSC or HGT models of gene tree discordance were used, DEPP was able to match the accuracy of maximum likelihood placement (and surpassed it when discordance was high) without any prior knowledge of the underlying model.

The model misspecification in our analyses came from gene tree discordance, but other forms of model misspecification exist and should be explored in future. Moreover, other reasons for discordance (e.g., the reference tree may be the taxonomic tree or built from morphological data) can also be imagined and provide potential use cases of DEPP. Finally, beyond accuracy, saving computational effort can provide a compelling reason to use black-box methods. If an expensive model is used to infer a tree, perhaps a black-box model can learn its essential features so that the tree can be extended further without repeating the effort. Such an approach would be presumably cheaper than applying the expensive model to the entire dataset. We leave the exploration of such applications to future work.

The specific formulation that we chose, embedding sequences in high dimensional spaces, allowed us to define a loss function that can be easily optimized using back propagation. One can argue that the ideal loss function for placement would be one that evaluates the accuracy of the final placement, not the distances. Designing such loss functions would be easy enough. However, optimizing a loss function with discrete components (e.g., the placement branch) will loose the differentiability of the loss function, which is necessary for backpropagation using standard methods. Finding ways to perform backpropagation in partially differentiable spaces like phylogenetic placement is an interesting topic, which we leave to future work.

Here, we trained our model on backbone trees that ranged in size from 200 to 10,000 species. While deep learning is believed to require extremely large labeled datasets for training, we were able to train DEPP, which has a moderate number of layers, with only 200 species because we use pairwise information. Thus, with 200 species, we have 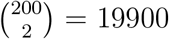 observed pairwise distances for training. Nevertheless, it is reasonable to expect that as reference trees become more densely sampled, the accuracy of DEPP would increase. Moreover, recall that our embedding is much smaller than size of the tree. Our results indicate that while theory suggests we need *n*−1 dimensions for Euclidean embedding of trees, far fewer dimensions suffice in practice; we had to reduce *k* from 128 to 32 to observe substantial drops in accuracy on simulated data (Table S2). Note that LR embedding states that *n* − 1 dimensions are sufficient, but it does not state that *n* − 1 dimensions are necessary. Future work should explore recent advances in hyperbolic neural networks (Ganea et al., 2018) and hyperbolic distances (Tabaghi and Dokmanić, 2020) to overcome limitations of Euclidean distances.

Our choices of hyper-parameters such as *k* and the training parameters such as the stopping criterion were based on pre-selected values that were not fine-tuned on any dataset. In extensive simulations, this simple procedure was necessary due to computational reasons. When applying in practice on real data, it is possible to fine-tune all the hyperparameters using a validation set. We can first randomly select a subset of the reference species as the validation set, use these as testing data to tune the parameters for the given dataset, and then train one last time with all the data with the fine-tuned parameters. Note that such a procedure will require repeated training and can be slow, and our preliminary results (Table S2) show that the gain in accuracy obtained can be small.

Our results on the real prokaryotic dataset indicate that DEPP *can* overcome horizontal transfer to find the right species tree placement, *as long as* similar patterns of HGT are observed in the training data. Note that HGT is the main cause of gene tree discordance in prokaryotes, even for the marker genes such as 16S (Gogarten et al., 2002; Stepanauskas et al., 2020). However, despite relatively low levels of error overall, a small tail of placements far away from the optimal species placement with errors *>*20 edges is observed for many genes (Fig. S8). This long tail may be a signature of horizontally transferred genes pointing to a very different position on the species tree than the genome-wide position. When HGT events are observed in the training data, DEPP has a chance to learn them and account for them. However, when an HGT event is novel (not seen among training data), DEPP has no way of recovering the correct position given just that one gene. These novel HGT events are a likely source of those infrequent cases of large error.

Similar to unobserved HGT events, unobserved sequence patterns can negatively impact the accuracy of DEPP. For the cases with high errors, we observe distances being overestimated, especially when the ML distances are low. We attribute this bias to the inability of the model to easily take advantage of novel data (e.g., changes in sites that are invariable in training data). We hope to remedy this limitation of our model in future work by more explicit modeling of unseen data, data augmentation, or changing the loss function.

The most immediate use of DEPP is in connecting 16S and metagenomics, as we demonstrated in our case study. DEPP often places 16S within three branches of the correct edge in the species tree; thus, while some errors remain, DEPP results enable combined 16S and metagenomic analyses with high accuracy. Currently, the main method used to combine results from 16S and metagenomic data is to use each data type to perform *taxonomic* identification and use the resulting classification in downstream analyses. Taxonomic classification clearly has less resolution than phylogenetic placement. DEPP (and more broadly, discordant placement) allows a phylogenetic, instead of taxonomic, approach for combining data. Once sequences from all the data types are added to the same tree, many downstream measurements, such as UniFrac distances and beta diversity, can be performed on the combined data. Our case study demonstrated that DEPP is capable of resolving real microbial community structures using either 16S amplicon or metagenomic data. Moreover, on this dataset, we observed improved ability to distinguish healthy and diseased samples by combining 16S and MAG data. Thus, combining both sources of data can reveal patterns that are relevant to the pathogenic profiles of the samples and the clinical status of the subjects. We saw remarkable levels of agreement between MAGs and ASV data in our case study, but also disagreements. While our data tended to provide evidence supporting the advantages of metagenomics over 16S amplicons, some limitations of MAGs (such as assembling similar species) were also revealed. Thus, due to pros and cons of each data type, the two sources of information are likely to remain complementary. Given 16S and metagenomic data, DEPP can add *all* samples to the same underlying tree, and this unified view of multiple data types enables downstream statistical analyses (e.g., Unifrac) to analyze both sets of samples jointly. By unifying the backbone tree, DEPP eliminates one source of bias, which is the use of separate reference libraries.

## Data and Code Availability

DEPP is publicly available at https://github.com/yueyujiang/DEPP.

Data from this paper are available at https://tera-trees.com/data/depp/.

## Supplementary Material

### A Supplementary Figures

**Figure S1:**
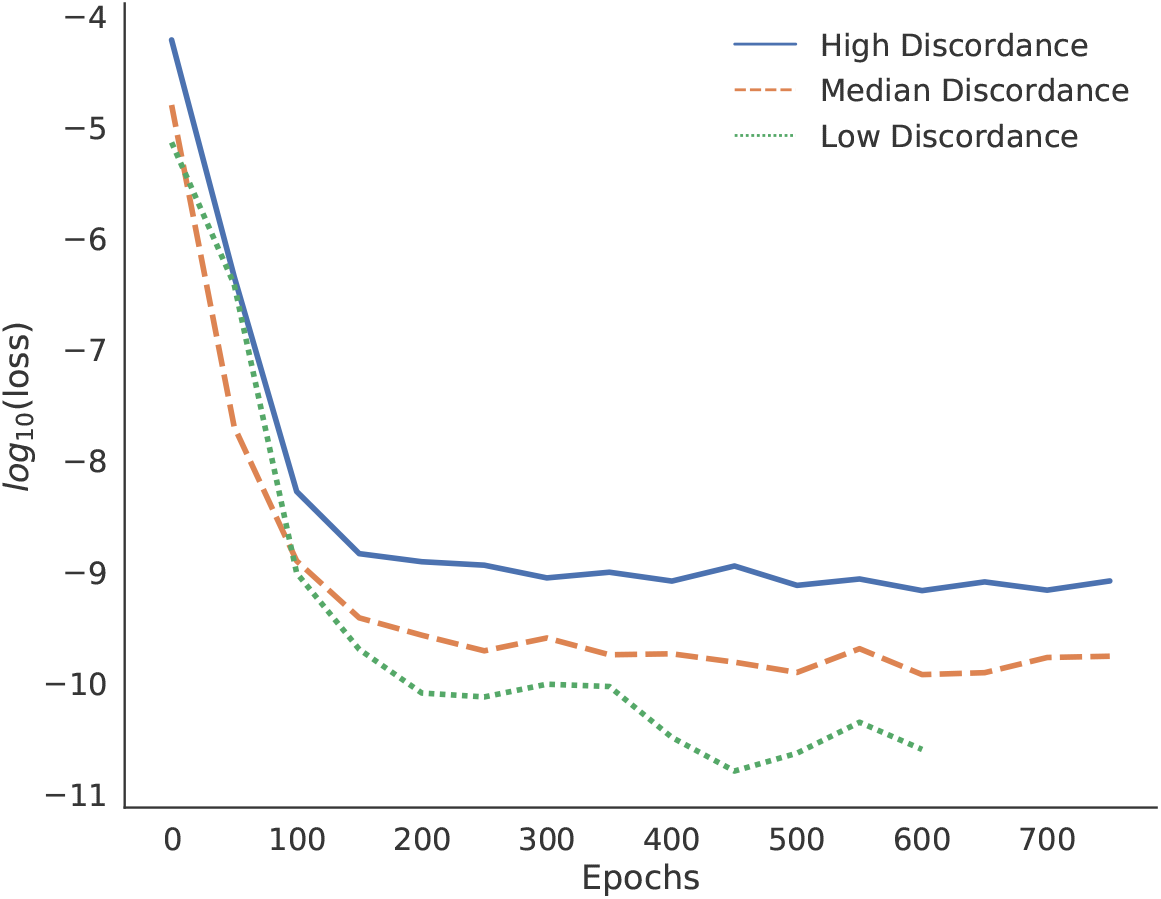
Train loss

**Figure S2:**
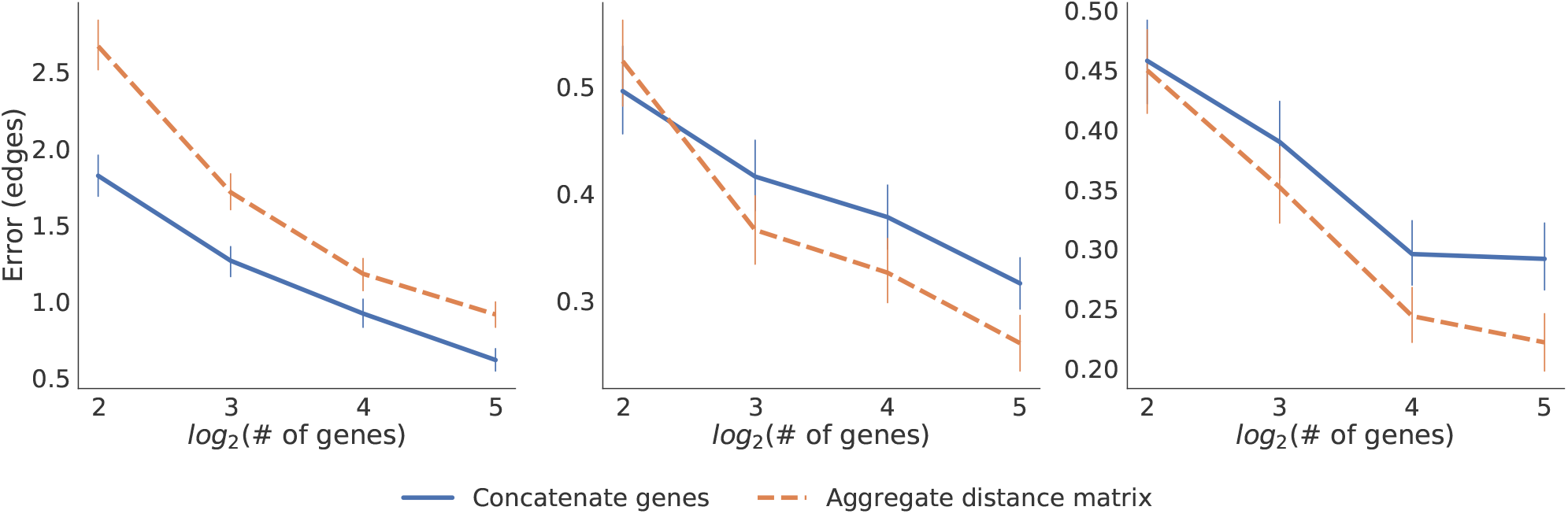
Mean and standard error of placement error versus the number of genes for simulated data. Blue: Results of single DEPP model trained by concatenating genes; Orange: Results of aggregating distance matrix from multiple DEPP models (one model per gene) by taking the mean distance among the interquartile range. Note that these analyses are using the older version of DEPP (v.0.1.56), and an older version of the dataset with ultrametric species trees used as the backbone (data release, v1.1.0).

**Figure S3:**
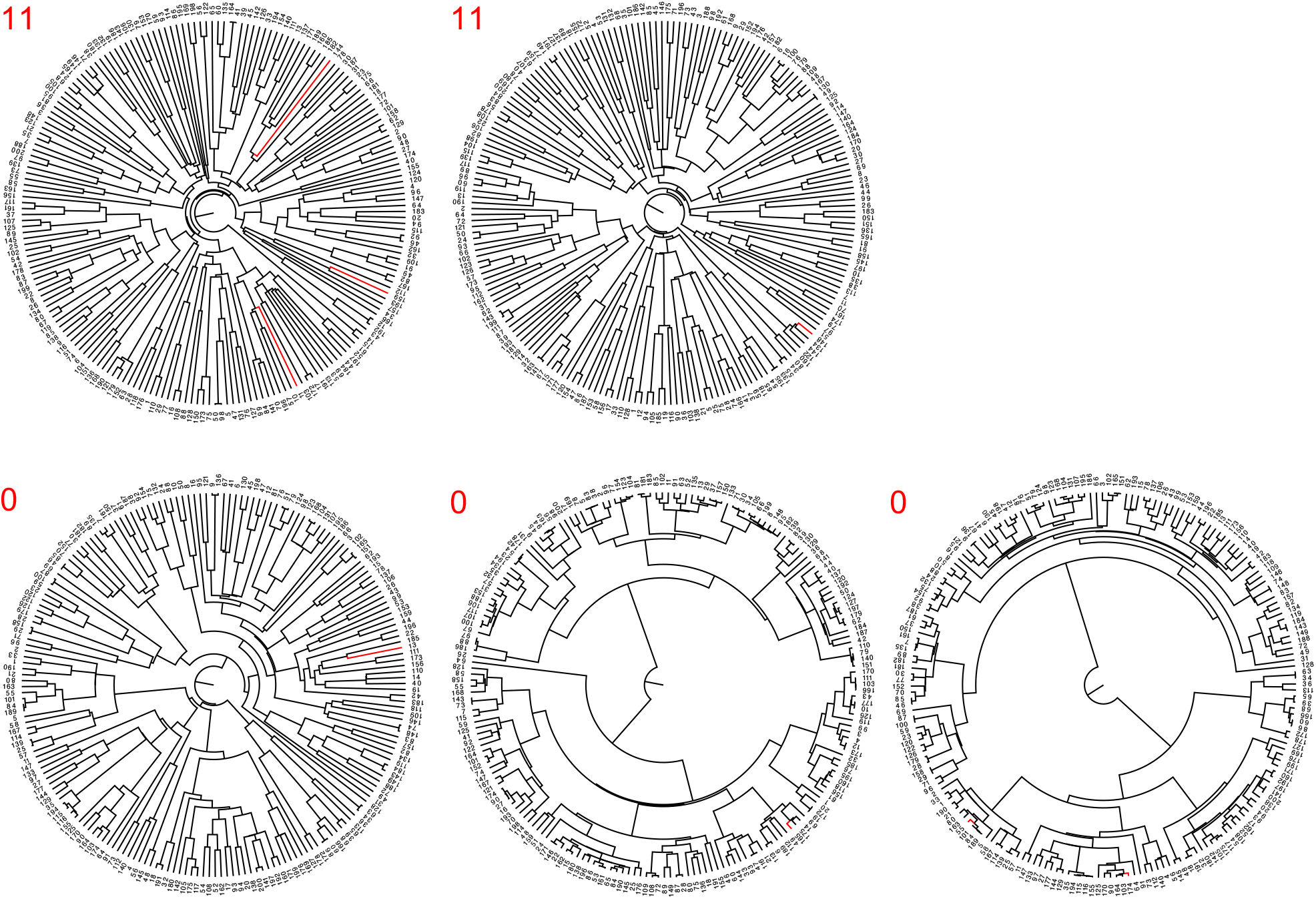
Examples of phylogenetic trees with high (top) and low (bottom) placement errors by DEPP. For each tree, we show the (correctly placed) query in red and show the error of DEPP on top in red.

**Figure S4:**
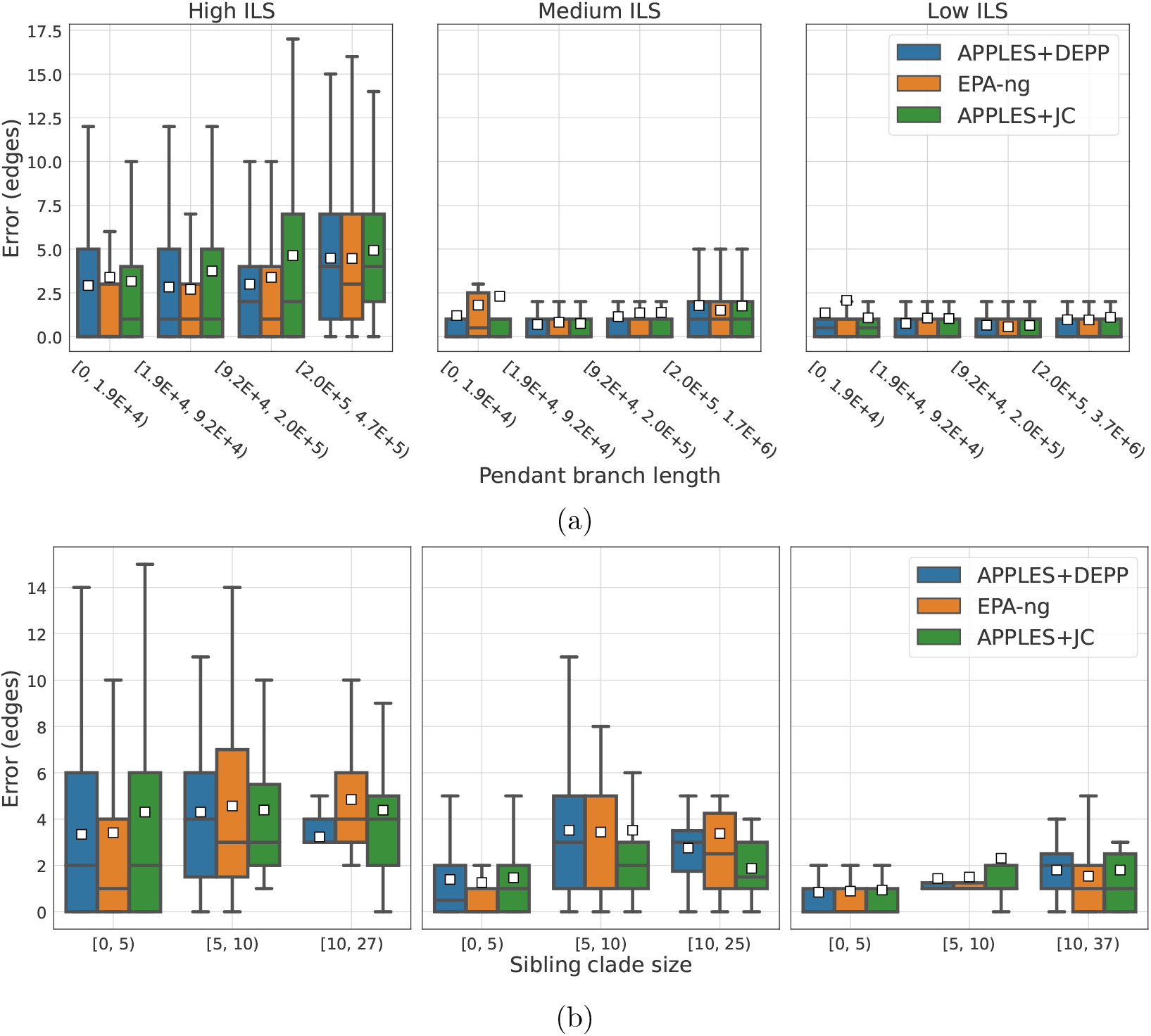
Impact of query properties on error. For each mode condition, we show the impact of the (a) length of the terminal branch of the query and (b) the number of leaves in the sibling clade of the query on the accuracy of all models when run with a single gene. The boxes correspond to levels of discordance, showing high, medium, and low from left to right.

**Figure S5:**
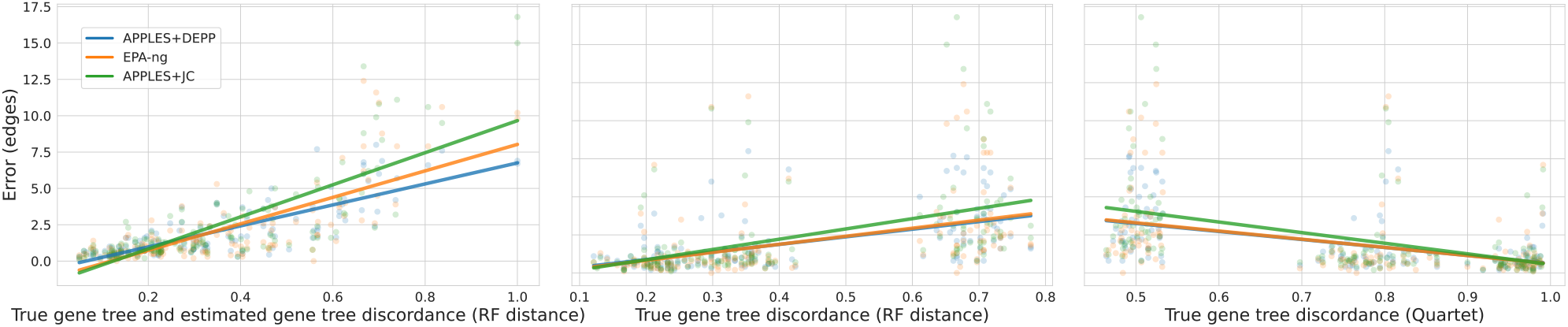
The impact of true gene tree discordance and lack of signal. left: Impact of lacking signal measured as the RF distance between true gene trees and the estimated gene trees. Middle: Impact of true gene tree discordance measured as the RF distance between true gene trees and the species tree. Right Impact of true gene tree discordance measured as the quartet score between true gene trees and the species tree, combining all discordance levels. Dots represent the mean of 10 queries for each of 148 replicate datasets.

**Figure S6:**
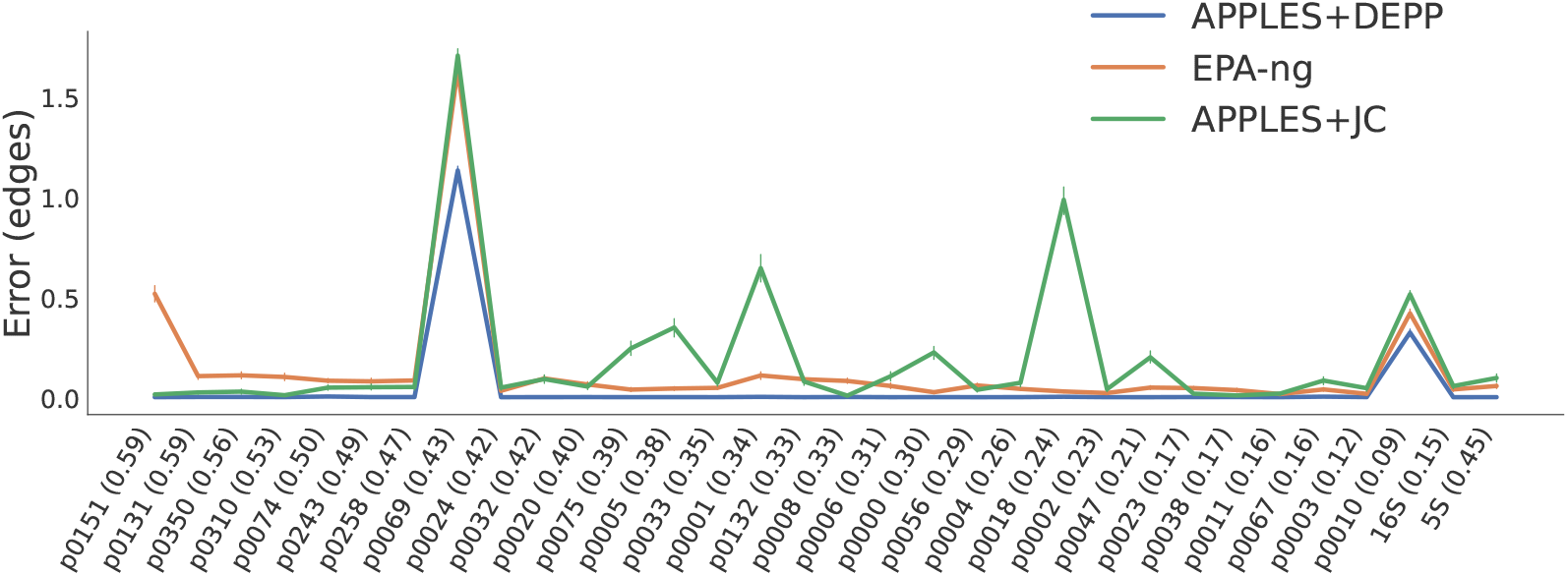
The mean and standard error of errors of APPLES+JC and EPA-ng versus DEPP when tested on known sequences (i.e., those in the reference set). Note that, unlike other genes, here, 16S and 5s have multiple copies (1.9 and 3.8 copies per species, respectively), causing the spike in the error.

**Figure S7:**
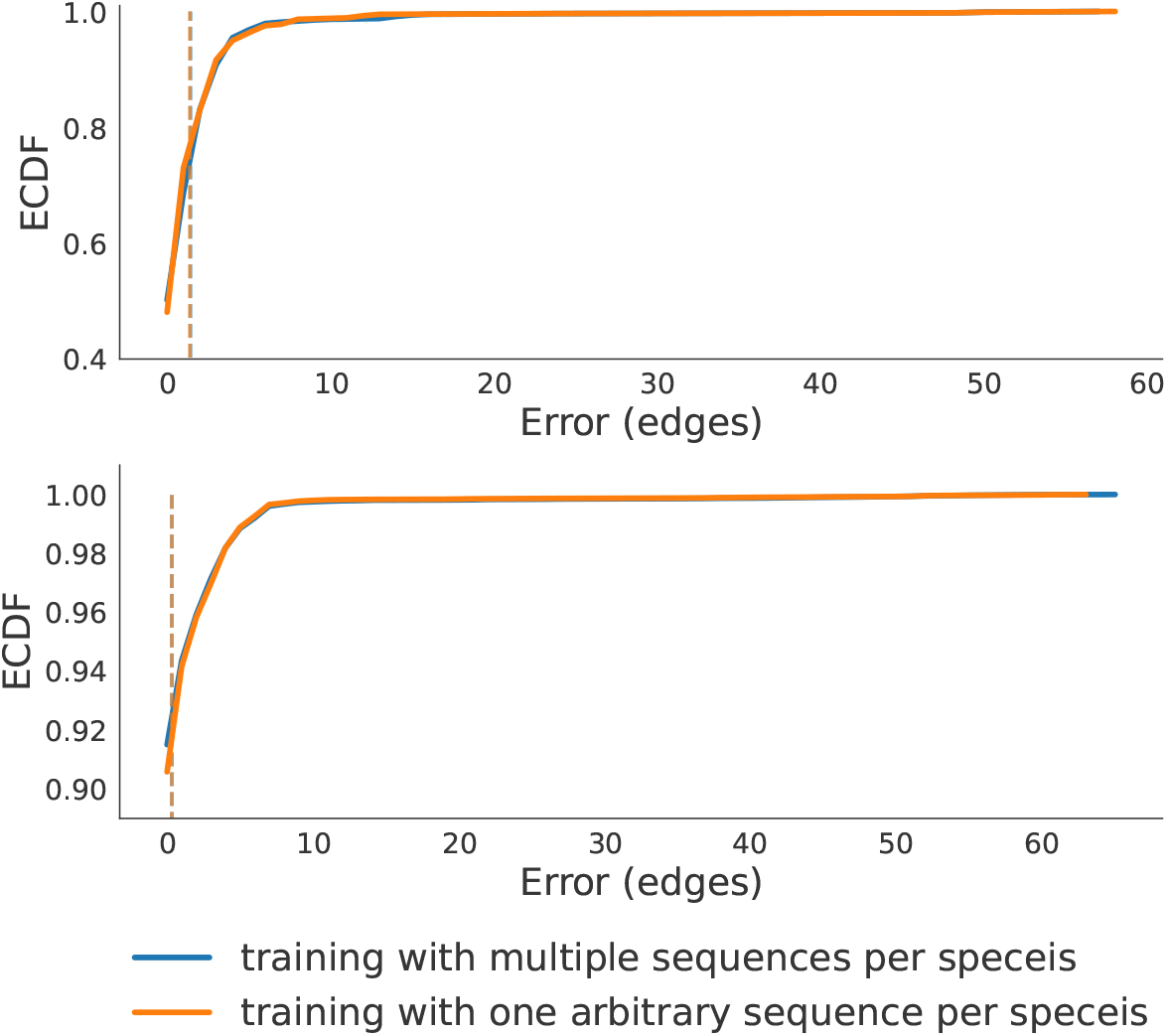
Impact of weighted encoding on WoL 16S data. The figure shows the comparison between training the model using all the available copies (using weighted encoding described in the text) versus selecting one arbitrary copy to represent each species. With multiple copies, sequenced are encoded by the frequency of the nucleotide character at each site. The results show that using more sequences per species has a very small impact on accuracy. Nevertheless, we use all the available sequences for training in all other experiments under the assumption that using all the sequences can reduce the variance of the DNN model. Results are based on version 1.0.0 of the DEPP WoL reference library.

**Figure S8:**
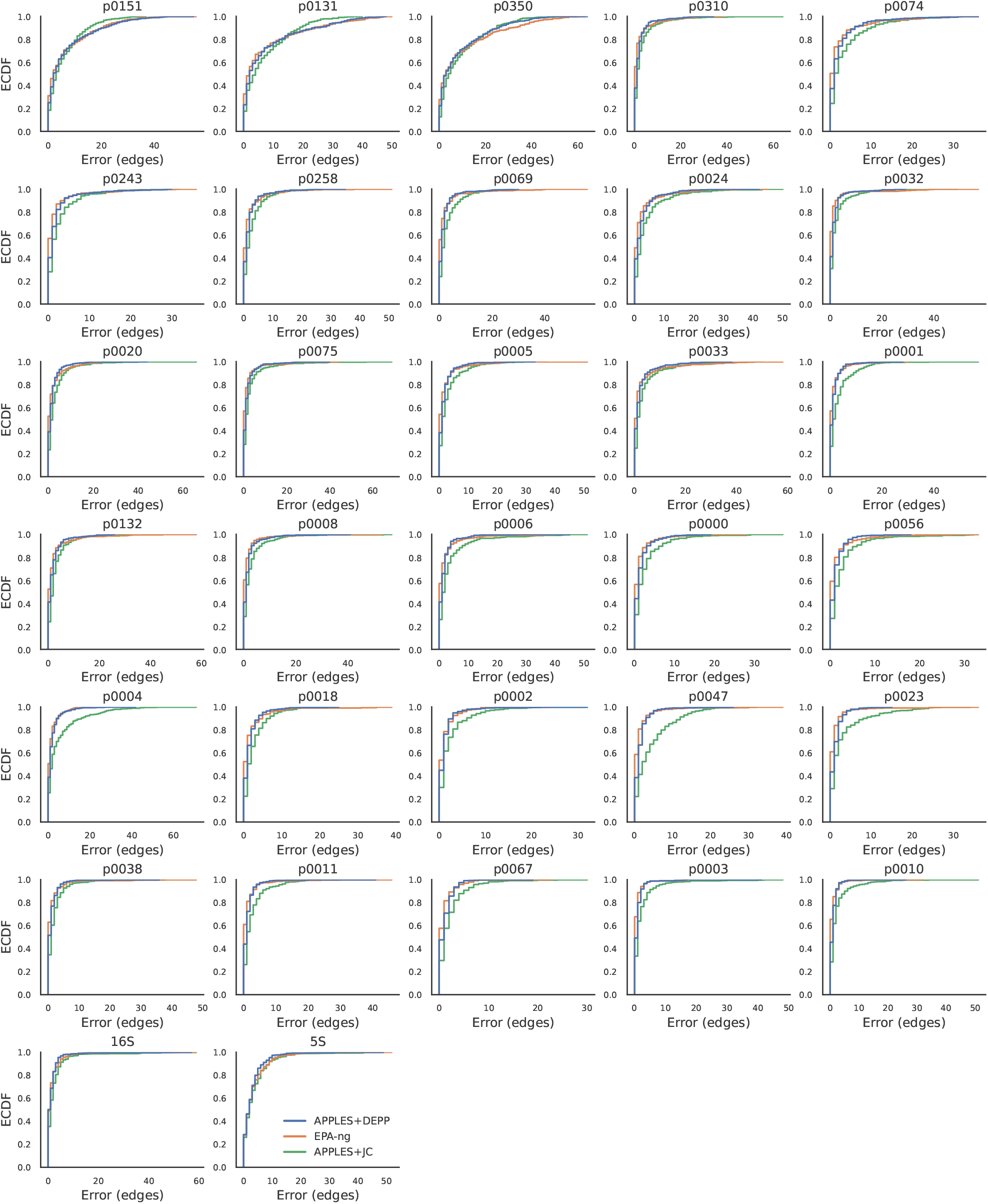
Placement error on real microbial data in leave-out-experiments. We show the Empirical Cumulative Distribution (EDCF) of the placement error for JC and DEPP per gene case. The top two rows correspond to genes with high discordance, the next two rows medium discordance, and the next two rows low discordance.

**Figure S9:**
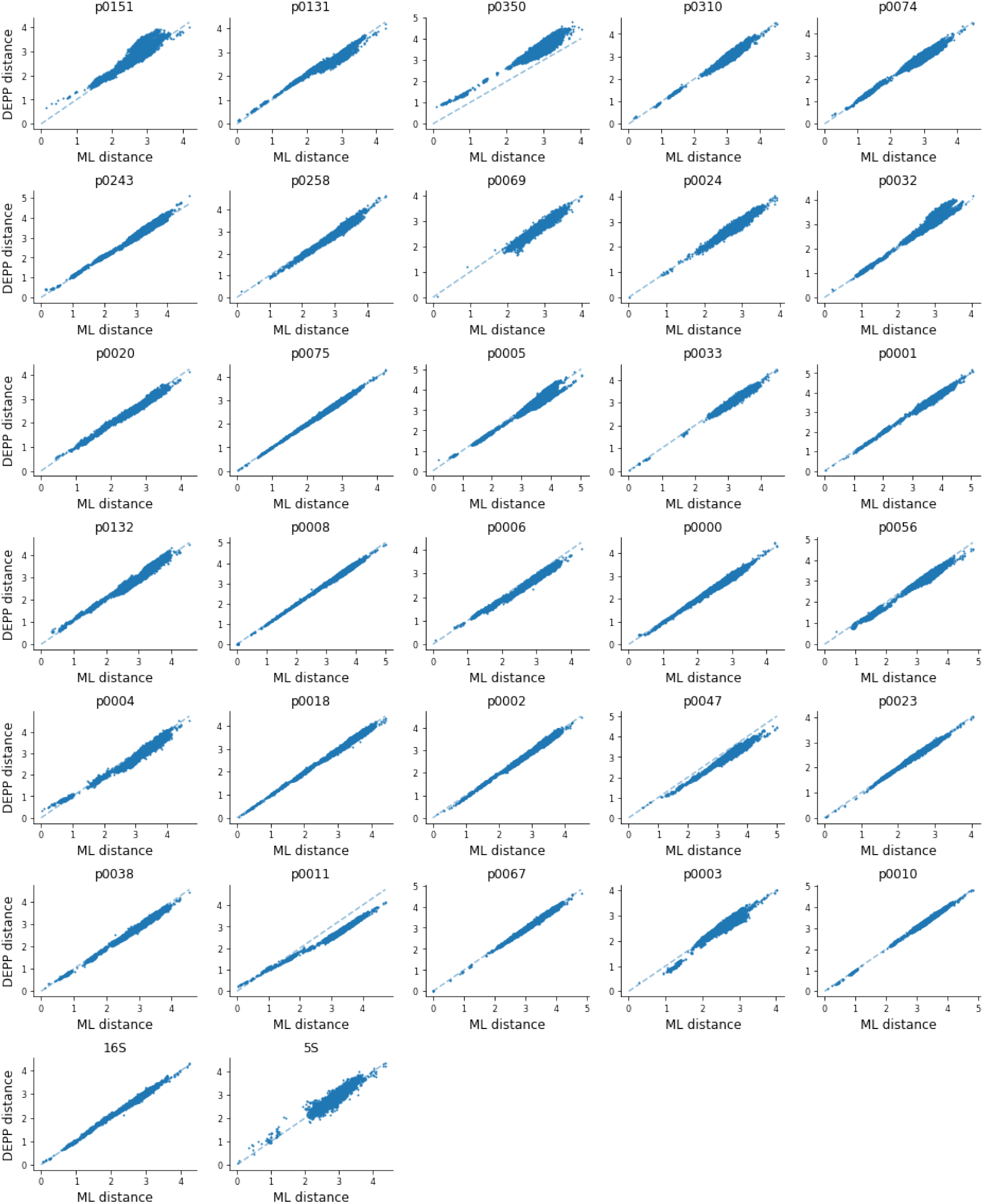
Examples of distance calculation using DEPP with errors equal to zero. x-axis: the true phylogenetic distance between the query; y-axis: distance calculated by DEPP. Each panel is a query arbitrarily chosen from the experiment corresponding to the gene indicated by the panel’s title. See Table S3 for gene properties.

**Figure S10:**
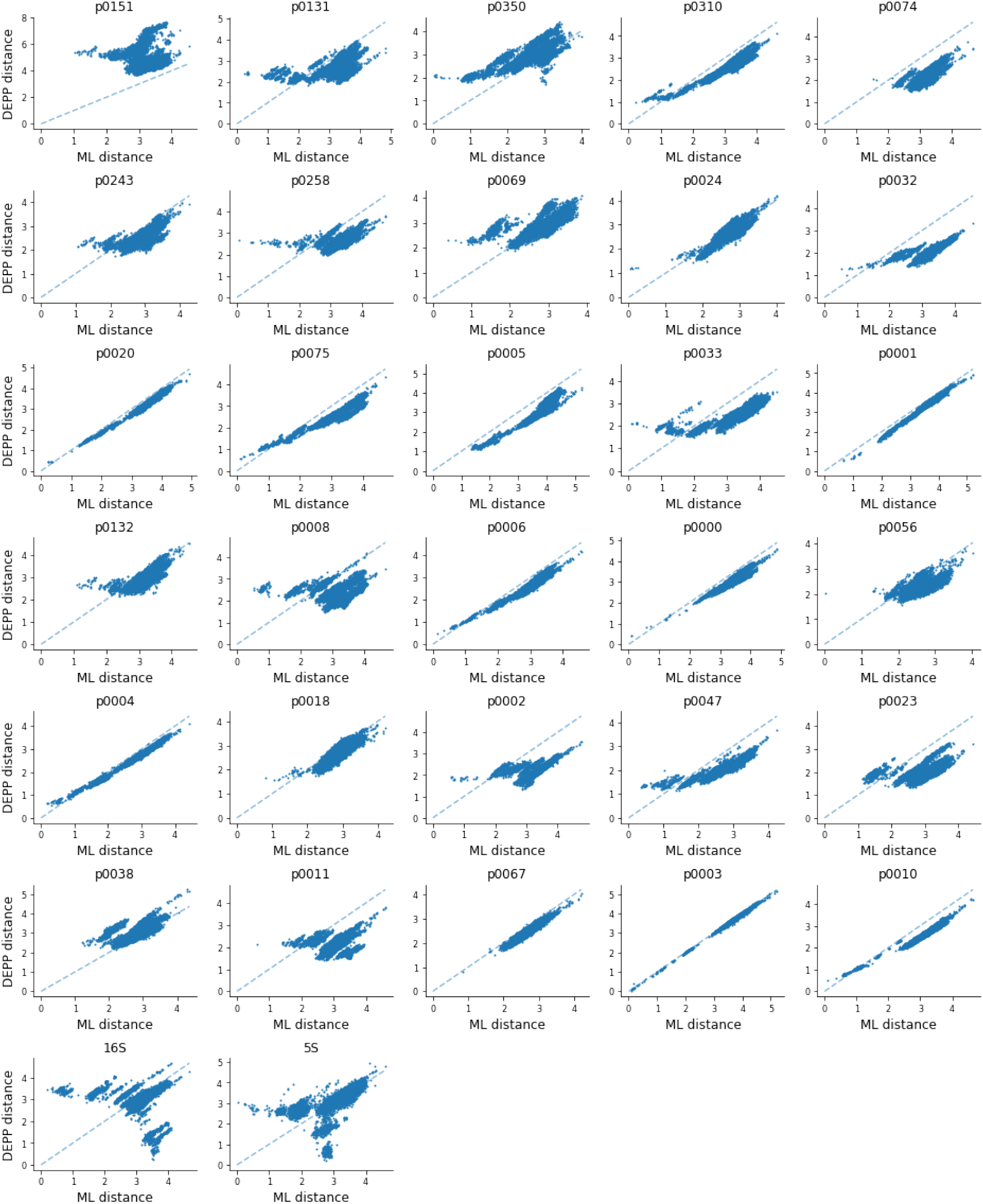
Examples of distance calculation using DEPP with errors larger than 15. x-axis: the true phylogenetic distance between the query; y-axis: distance calculated by DEPP. Each panel is a query arbitrarily chosen from the experiment corresponding to the gene indicated by the panel’s tile. See Table S3 for gene properties.

**Figure S11:**
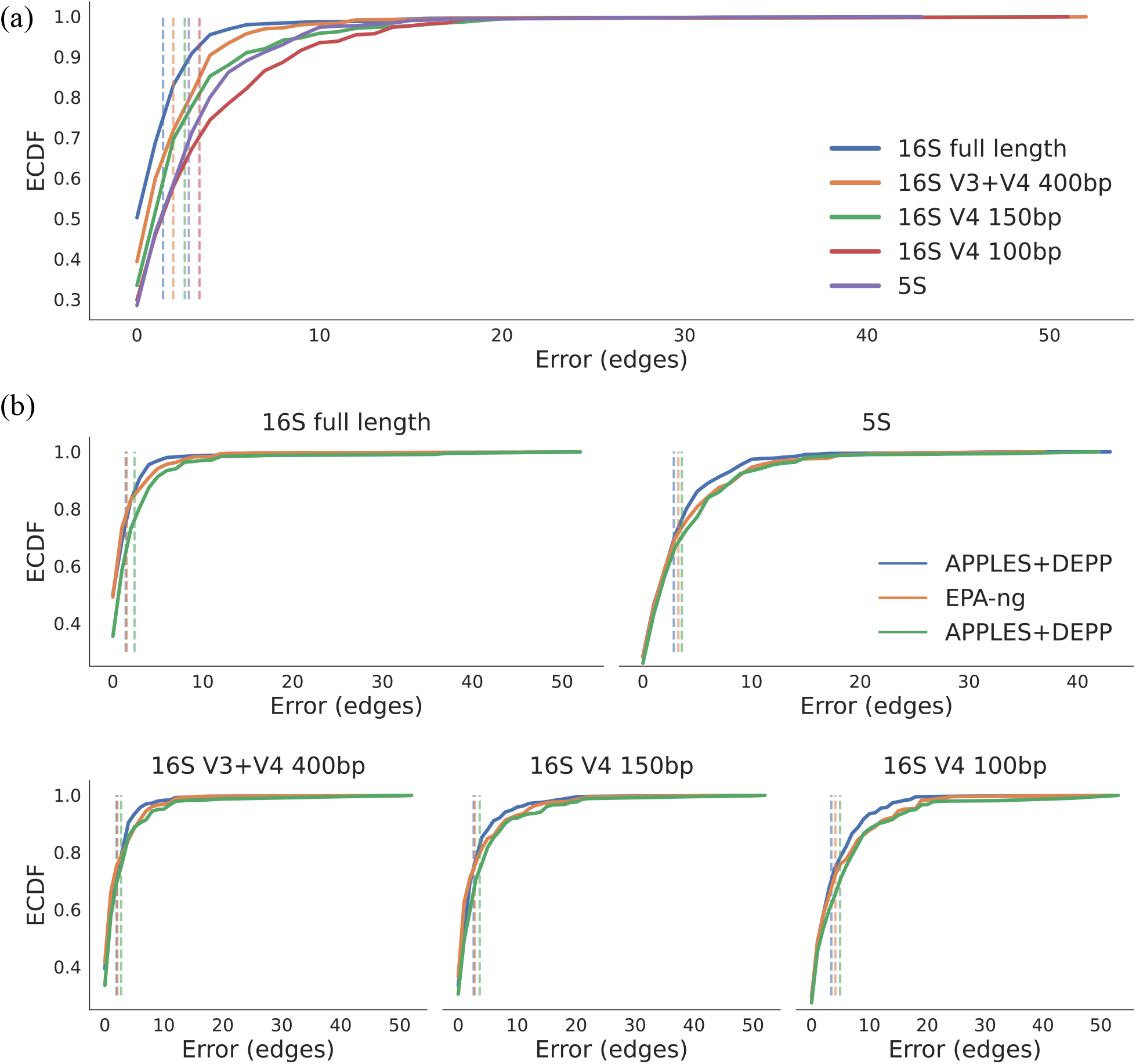
Complete Empirical CDF on the WoL dataset. In this figure we show the complete ECDF figure on WoL data; a truncated version is shown Fig. 4b.

**Figure S12:**
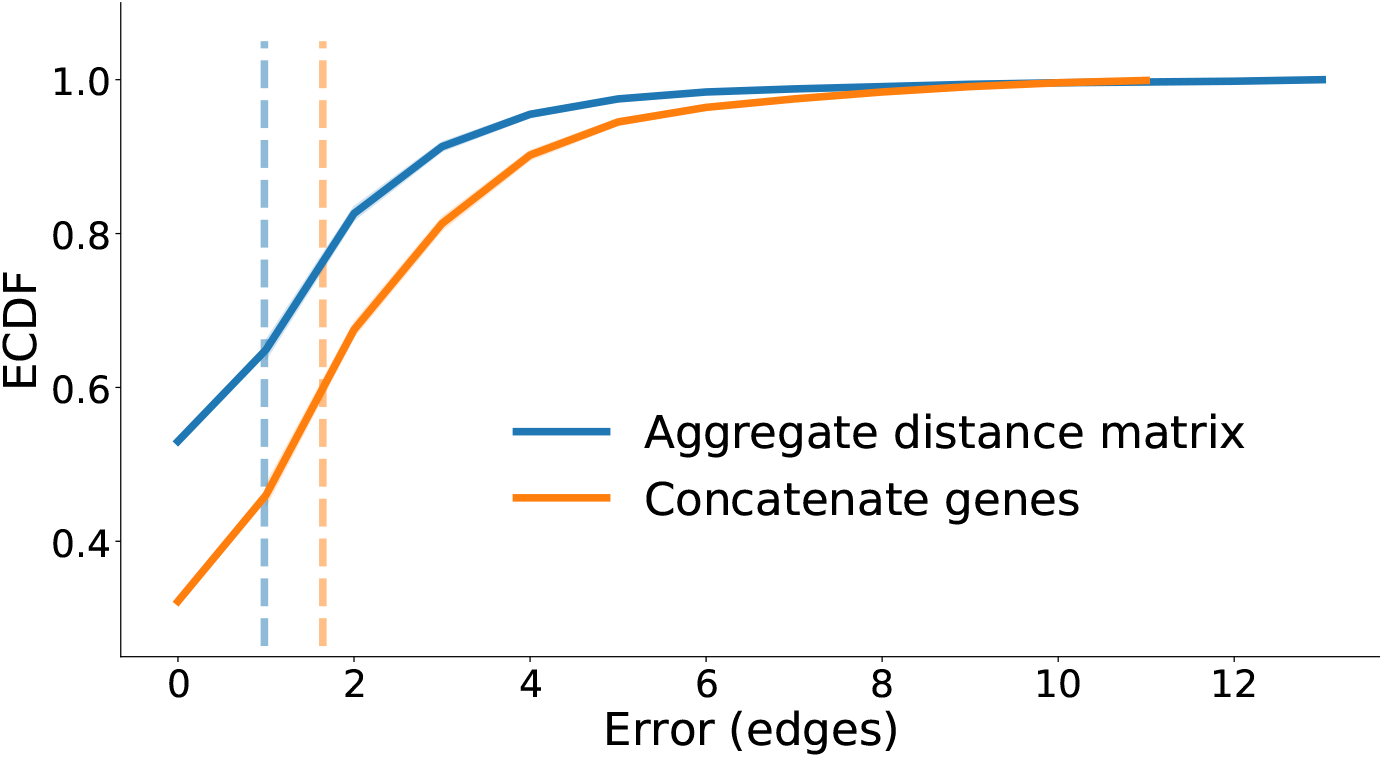
Results of multiple genes on WoL dataset summarized using two different ways. We show results of using 50 randomly selected genes in a leave-out experiment. Blue: We train 50 models which gives us 50 distance matrices. We then aggregate the 50 distance matrices by taking the mean distances in the interquartile range. Orange: We concatenate the sequences from 50 genes and train a model on the concatenated sequences. Note that these analyses are using the older version of DEPP (v.0.1.56), and an older version of the dataset with ultrametric species trees used as the backbone (data release, v1.1.0).

**Figure S13:**
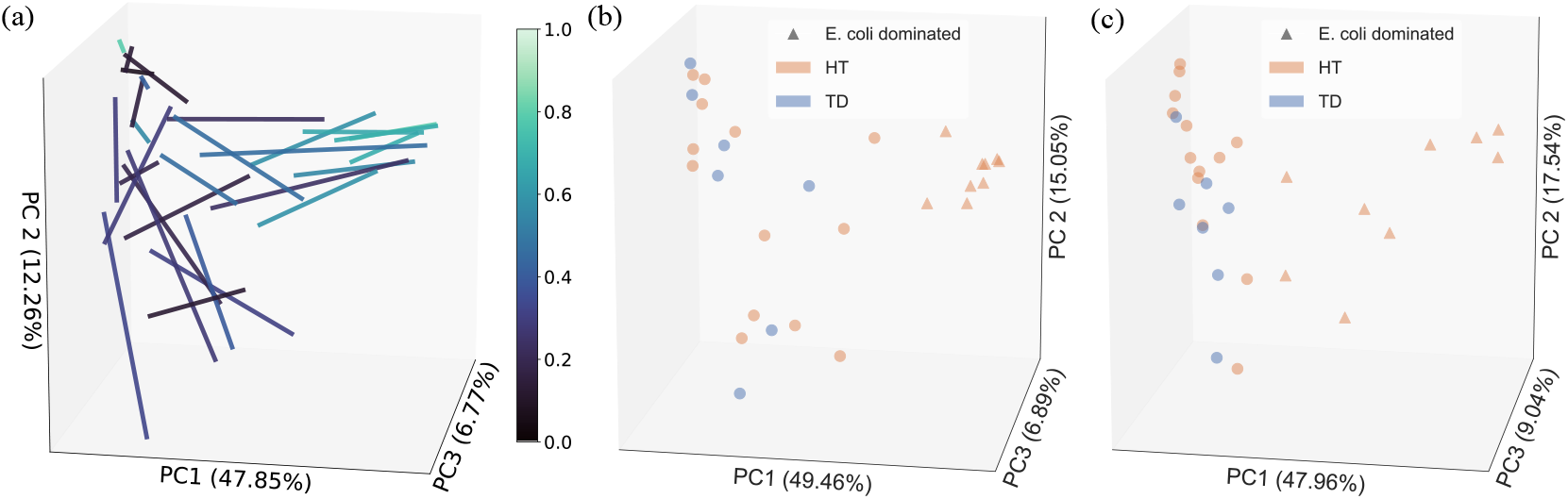
Visualization of beta diversity of Traveler’s Diarrhea (TD) gut microbiomes. We plot samples using PCoA run on weighted UniFrac distances. a) The PCoA visualization of both MAG and ASV profiles. Profiles from the same sample are linked by an edge, and the color of the edge shows the percentage of sequencing data covered by MAGs. Note that lighter edges, corresponding to samples with a higher MAG coverage, tend to have shorter lengths. b,c) The PCoA visualization of MAG profiles (b) and ASV profiles (c) separately. *E. coli*-dominated samples are more easily delineated using MAGs.

**Figure S14:**
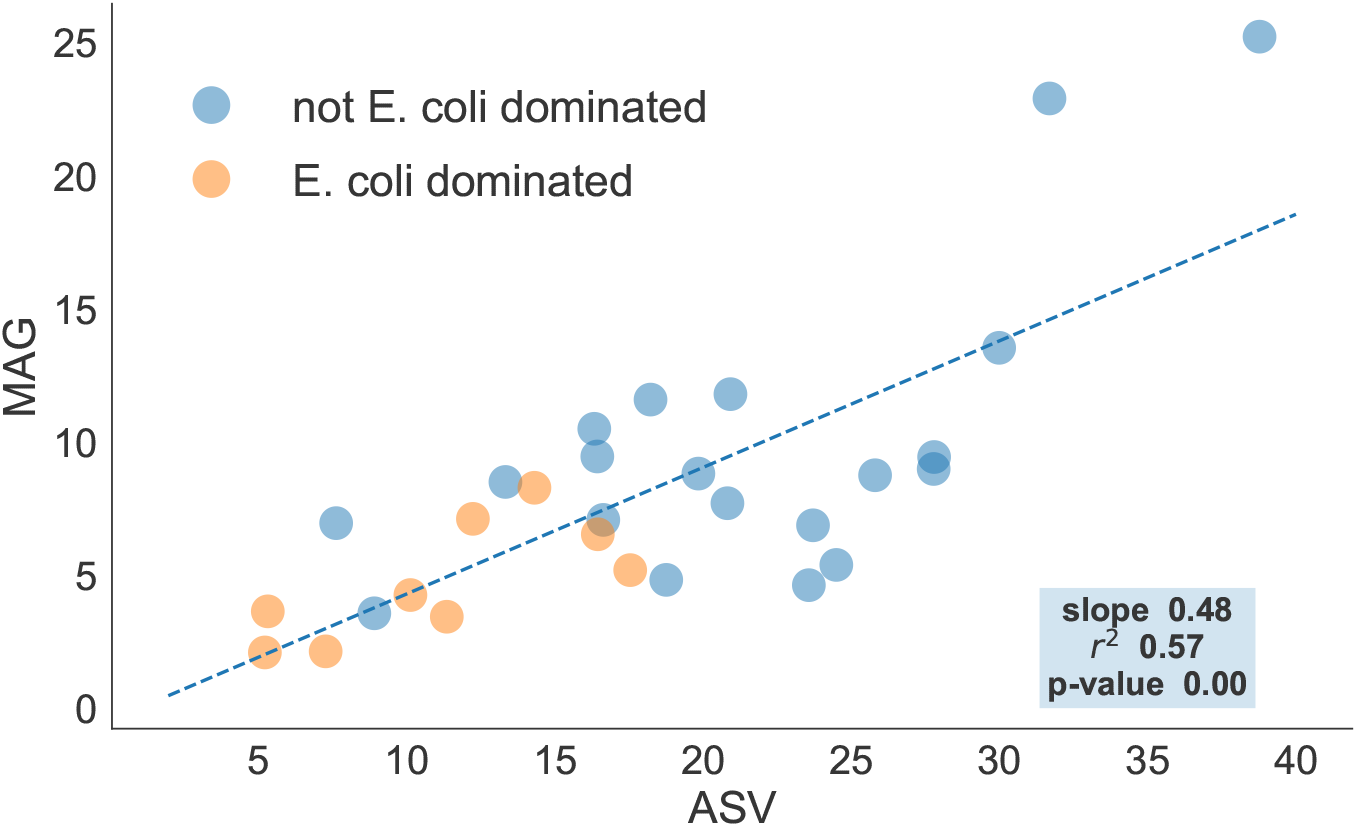
Correlation of Alpha diversity measured using ASVs and MAGs on Traveler’s Diarrhea dataset. We show the alpha diversity of MAGs versus ASVs, measured using Faith’s phylogenetic diversity (PD). Since this measure ignores abundance, to remove the impact of noise, we only consider the most abundant species that cumulatively cover at least 95% of the placed sequences for each sample. The dotted line shown in the figure is fitted using alpha diversity from all 29 samples.

**Figure S15:**
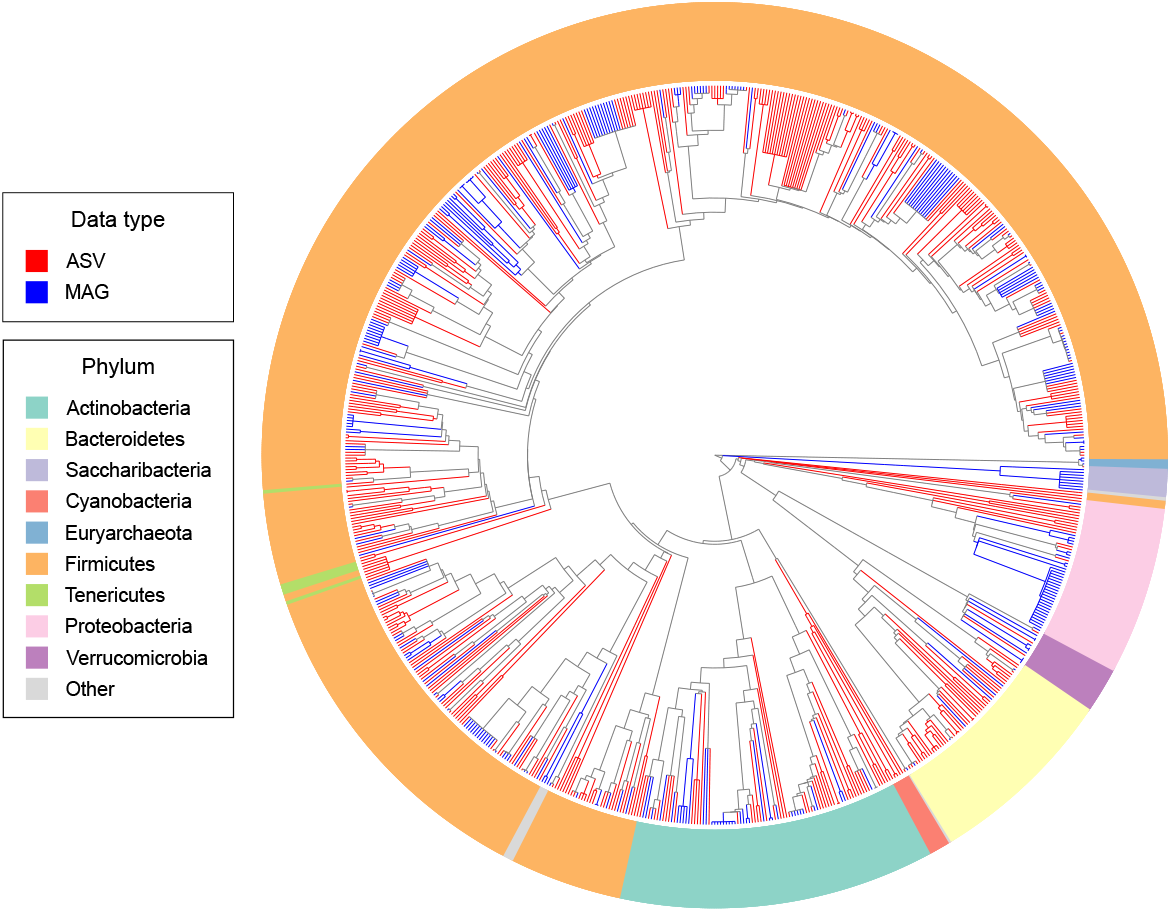
ASV and MAG placement on the WoL phylogenetic tree (no backbone species included)

**Figure S16:**
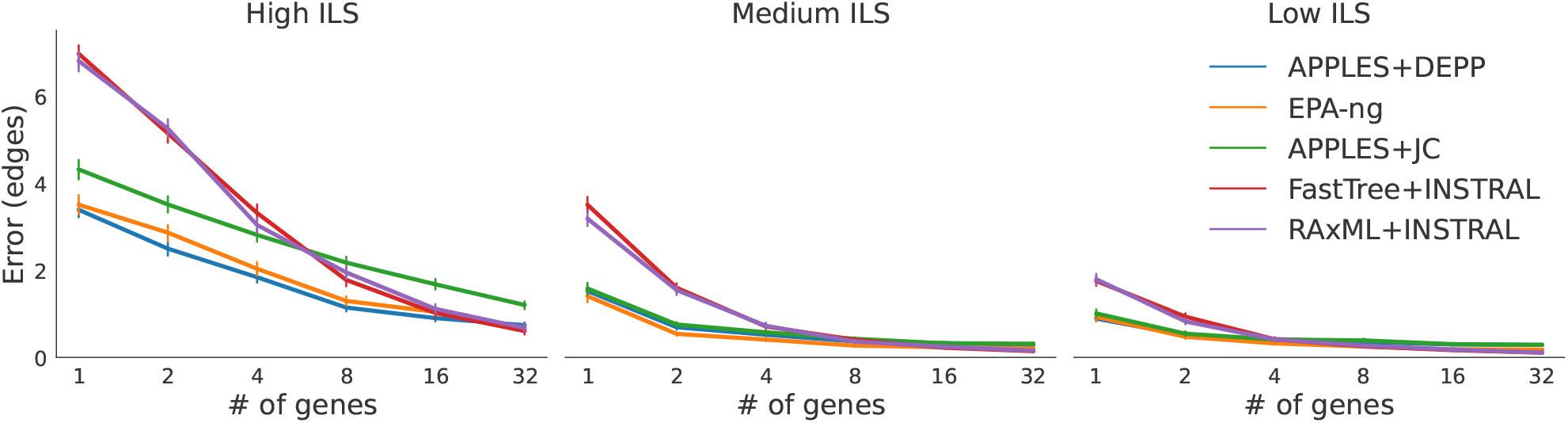
Mean and standard error of placement error versus the number of genes. Input tree to INSTRAL is estimated using FastTree and RAxML

**Figure S17:**
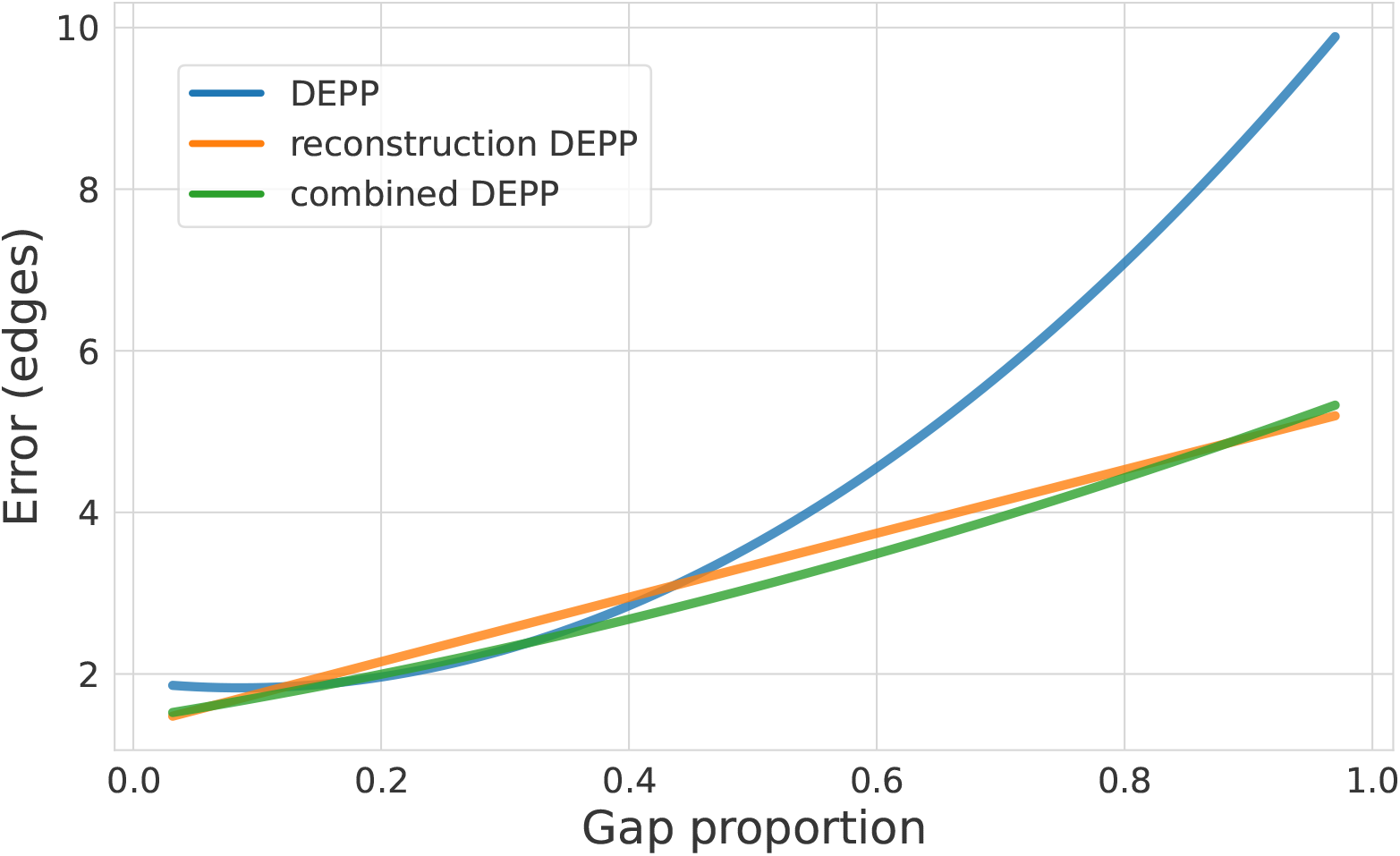
Impact of reconstruction network. The x-axis is the proportion of gaps in the alignments of the queries; Data is from 30 marker genes in the WoL dataset

### B Supplementary Tables

**Table S1:**
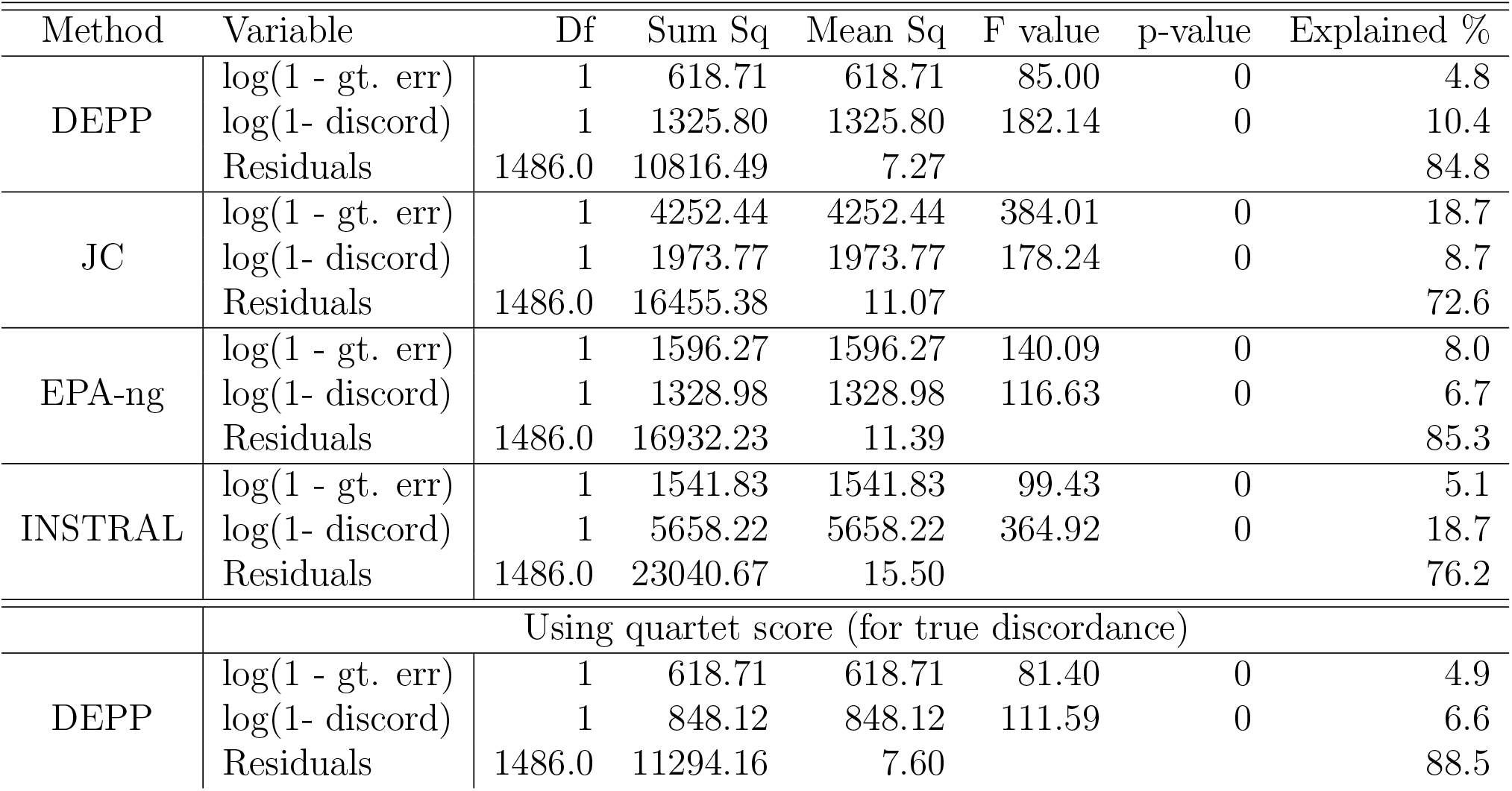
ANOVA statistical tests of the impact of gene tree discordance (RF between true gene trees and the species tree) and lack of gene signal (RF between true gene trees and estimated gene trees), both log transformed for better linearization, on the placement error of DEPP and APPLES+JC.

| Method | Variable | Df | Sum Sq | Mean Sq | F value | p-value | Explained % |
| --- | --- | --- | --- | --- | --- | --- | --- |
| DEPP | log(1 - gt. err) | 1 | 618.71 | 618.71 | 85.00 | 0 | 4.8 |
|  | log(1- discord) | 1 | 1325.80 | 1325.80 | 182.14 | 0 | 10.4 |
|  | Residuals | 1486.0 | 10816.49 | 7.27 |  |  | 84.8 |
| JC | log(1 - gt. err) | 1 | 4252.44 | 4252.44 | 384.01 | 0 | 18.7 |
|  | log(1- discord) | 1 | 1973.77 | 1973.77 | 178.24 | 0 | 8.7 |
|  | Residuals | 1486.0 | 16455.38 | 11.07 |  |  | 72.6 |
| EPA-ng | log(1 - gt. err) | 1 | 1596.27 | 1596.27 | 140.09 | 0 | 8.0 |
|  | log(1- discord) | 1 | 1328.98 | 1328.98 | 116.63 | 0 | 6.7 |
|  | Residuals | 1486.0 | 16932.23 | 11.39 |  |  | 85.3 |
| INSTRAL | log(1 - gt. err) | 1 | 1541.83 | 1541.83 | 99.43 | 0 | 5.1 |
|  | log(1- discord) | 1 | 5658.22 | 5658.22 | 364.92 | 0 | 18.7 |
|  | Residuals | 1486.0 | 23040.67 | 15.50 |  |  | 76.2 |
| Using quartet score (for true discordance) |  |  |  |  |  |  |  |
| DEPP | log(1 - gt. err) | 1 | 618.71 | 618.71 | 81.40 | 0 | 4.9 |
|  | log(1- discord) | 1 | 848.12 | 848.12 | 111.59 | 0 | 6.6 |
|  | Residuals | 1486.0 | 11294.16 | 7.60 |  |  | 88.5 |

**Table S2:** Impact of hyperparameters on the error of DEPP. We test on the single-gene high discordance 200-taxon simulated dataset. Bold: the default setup, used for the results reported in the paper. *k*: embedding size. To change the architecture, we change the number of resblocks. Various weighting schemes are incorporated in (1). Error: mean placement error (edges). Statistical tests of significance compare each condition to the default condition using a two-sided paired Student’s t test (*p*-values).

|  | 32 | 64 | 128 | 128 | **128** | 128 | 128 | 128 | 256 |
| --- | --- | --- | --- | --- | --- | --- | --- | --- | --- |
| resblocks | 5 | 5 | 1 | 3 | **5** | 5 | 5 | 5 | 5 |
| weighting | $\frac{1}{\delta^2}$ | $\frac{1}{\delta^2}$ | $\frac{1}{\delta^2}$ | $\frac{1}{\delta^2}$ | $\frac{1}{\delta^2}$ | 1 | $\frac{1}{\delta}$ | $\frac{1}{\delta^4}$ | $\frac{1}{\delta^2}$ |
| error | 3.836 | 3.380 | 3.734 | 3.534 | **3.442** | 3.374 | 3.516 | 3.520 | 3.302 |
| $p$-values | 0.002 | 0.544 | 0.023 | 0.359 | N/A | 0.472 | 0.454 | 0.442 | 0.161 |

*Observations:*
- Increasing embedding size ( $k$ ) tend to reduce the error. - More residual blocks lead to better performance. - DEPP is robust to weighting schema. Different weighting schema do not have statistically significant difference in error.

**Table S3:**
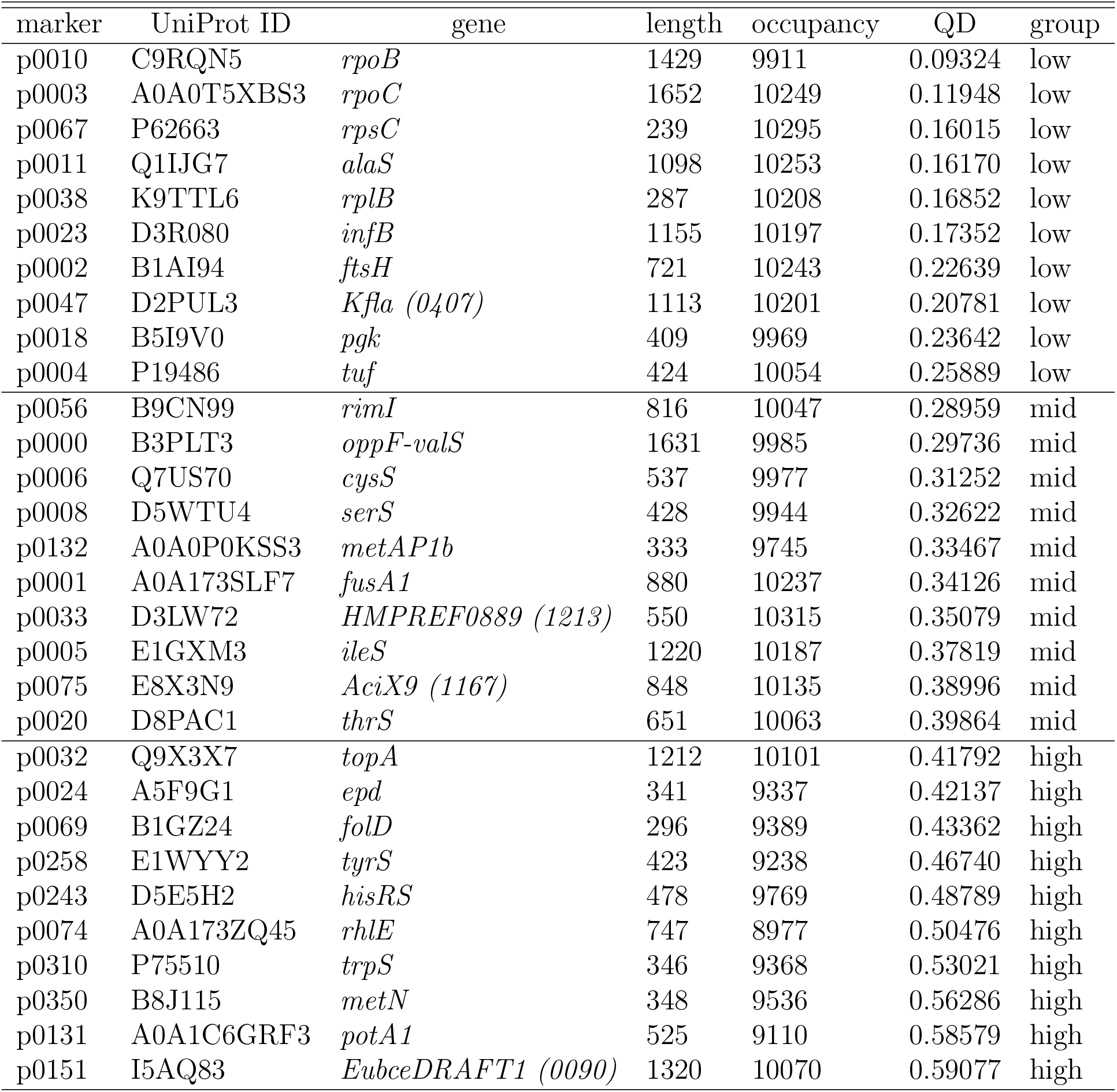
Properties of selected genes from the WoL dataset Zhu, Mai, et al., 2019. Length indicates the length of the gene’s protein alignment. Occupancy shows the number of species that has the each gene in their genome. QD: Quartet distance between the published gene tree and the species tree. Finally, group shows the assignment of genes to the three groups of low, medium, and high discordance based on the QD.

**Table S4:**
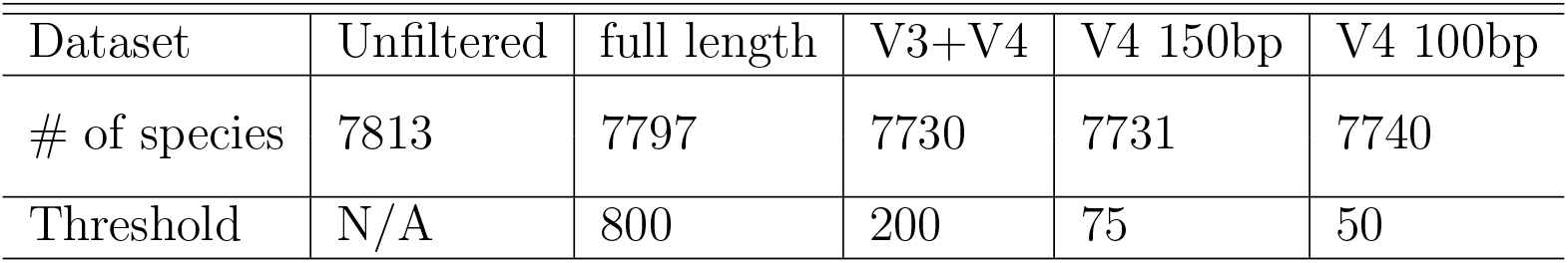
Number of species left after filtering out sequences that is shorten than roughly half the average length.

### C Background on CNNs

#### Convolutional layers

Denote an input feature map of the intermediate layer of a model as 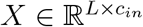, where *L* is the length of the feature map (here, sequence length), and *c*_*in*_ is the number of input channels (here, 4 for the initial layer). Also given is a filter 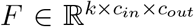 where *k* is the kernel size and *c*_*out*_ is the number of output channels. The convolutional operation is defined as:

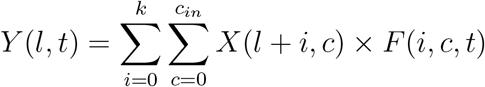

One-dimensional convolutional layer use sliding windows to process the input sequence. Each site in the output is the weighted sum of the fragment in the corresponding windows plus the bias.

In a neural network, convolutional layers are usually used as feature extractors. Multiple convolutional layers are able to detect high-level abstraction from the input. One-dimensional convolutional layer use sliding windows to process the input sequence. Each site in the output is the weighted sum of the fragment in the corresponding windows plus the bias. The weights are determined by a learnable kernel.

#### Fully Connected Layer

In a neural network, fully connected layers are usually used as the last few layers to aggregate information and get the final output. In the fully connected layer, each activation in the output has connections with all the input. Specifically, each output dimension is a weighted sum of all the input, and the weights are optimized during training.

#### Nonlinear layer

Convolutional layers and fully connected layers consist of only linear operations. Introducing nonlinear layers can give the model the ability to capture nonlinear relations. While many nonlinear kernels exist, in this work, we use Continuously Differentiable Exponential Linear Units (CELU) as the nonlinear layer applied at the end of each layer. CELU has the following form:

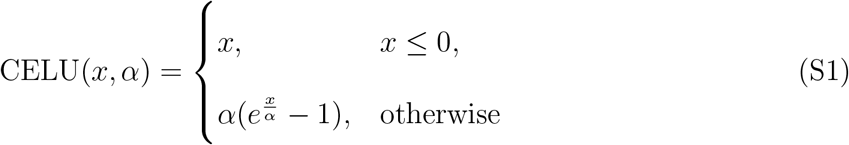

### D Exact commands and versions

#### Software version

- APPLES 2.0.0
- DEPP 0.2.2
- EPA-ng 0.3.8
- INSTRAL 5.13.5
- RAxML-ng 1.0.1
- RAxML 8.2.12
- UPP 4.3.10
- Qiime2 2020.11.1
- Prodigal v2.6.3
- PhyloPhlAn commit 2c0e61a
- Simphy 1.0
- Indelible V1.03

#### Branch length reestimation

- `raxml-ng --evaluate --msa $sequence file --tree $tree --model JC --threads 2 --blopt nr_safe`
- `raxmlHPC-PTHREADS -s $sequence file -w $outdir -n run -p 12345 -T 32 -m GTRCAT -g $tree`

#### Detecting Genes

- Predict open reading frames (ORFs) and translate DNA sequences into amino acid sequences
  – `prodigal -p $mode -f gff -g 11 -i $g.fna -o $g.gff -a $g.faa -d $g.ffn` `$g`: genome ID, $`mode`: “single” for single genomes (e.g., WoL) and “meta” for metagenomes (e.g., TD)
- Identify the 400 marker genes from ORFs
  – `phylophlan.py` `--c_dat $tmpdir –c_in $indir` `--c_out $outdir --nproc $cpus -u $rundir`

#### Gene alignment

Genes were aligned using UPP using the following commands for aligning the backbone sequences(first command), and aligning the novel queries to the existing backbone alignments (second command).

- `Run_upp.py -s $input_seq -B 2000 -M -1 -T 0.33 -A 200`
- `Run_upp.py -s $query_seq -a $backbone_seq -t $backbone_tree -A 100 -d $outdir`

#### Placement

- APPLES+DEPP
  – Training
    * Simulated dataset `Train_depp.py` `Backbone_tree_file=`$`backbone_tree` `Backbone_seq_file=`$`backbone_seq` `patience=5 lr=1e-4 embedding_size=128`
    * Marker genes in WoL dataset `Train_depp.py` `backbone_tree_file=$backbone_tree` `backbone_seq_file=$backbone_seq` `patience=5 lr=1e-4 embedding_size=512`
    * 5S/16S in WoL dataset `Train_depp.py` `Backbone_tree_file=$backbone_tree` `Backbone_seq_file=$backbone_seq` `patience=5 lr=1e-4 embedding_size=512 replicate_seq=True`
    * Note: for version 1.0.0 of this reference DB, we used DEPP 0.1.13, for version 1.1.0 (used everywhere unless otherwise specified), we use DEPP 0.1.54 for version 1.2.0, we use DEPP 0.2.2.
  – Query time
    * Calculating distance matrix
      · `Depp_distance.py` `Query_seq_file=$query_seq` `backbone_seq_file=$backbone_seq` `model_path=$model_path` `outdir=$out_dir`
    * Placement using APPLES
      · `run apples.py -d $distance_file -t $backbone_tree -o -f 0 -b 5`
- APPLES+JC `run apples.py -s $backbone tree -q $query seq -t $backbone_tree -f 0 -b 5`
- EPA-ng
  – `raxml-ng` `--evaluate` `--msa $backbone_seq` `--tree $backbone_tree` `--prefix info` `--model GTR+G+F` `--threads 2 --blopt nr_safe`
  – `epa-ng` `--ref-msa $backbone_seq` `--tree $backbone_tree` `--query $query_seq` `--model $GTR_info`
- INSTRAL
  – `java -Djava.library.path=./ -jar __instral.jar__` `-i $gene_trees` `-f $backbone_tree` `--placement $query` `-o $placement tree -C`

#### Weighted UniFrac

- `qiime diversity beta-phylogenetic` `--p-metric weighted_unifrac` `--i-table $feature_table` `--i-phylogeny $placement_tree` `--o-distance-matrix $unifrac_distance matrix`

#### PERMANOVA

- `qiime diversity beta-group-significance` `--i-distance-matrix $unifrac_distance_matrix` `--m-metadata-file $metadata` `--o-visualization $permanova_output` `--m-metadata-column group` `--p-permutations 999999`

#### PCoA

- `qiime diversity pcoa` `--i-distance-matrix $unifrac distance matrix --o-pcoa $pcoa_matrix`

#### D.1 HGT simulation

- `simphy -rs 10 -rl f:500 -rg 1 -sb f:0.0000005 -sd f:0.000000416666667` `-st f:100000000 -sl f:10000 -si f:1 -sp f:500000 -su f:4e-08 -hh f:1` `-hs ln:1.5,1 -hl ln:1.3692114,0.6931472 -hg ln:1.5,1 -cs 14907 -gt n:-18,0.4` `-lt ln:gt,0.75 -lk 1 -lb f:0 -ld f:0 -v 3` `-o model.10000.100000000.0.0000005.norm-lognormal -ot 0 -op 1 -od 1 > log.txt`

